# *Achromobacter xylosoxidans* isolates exhibit genome diversity, variable virulence, high levels of antibiotic resistance and potential intrahost evolution

**DOI:** 10.1101/2025.07.29.667465

**Authors:** Pooja Acharya, Cameron Lloyd, Ngoc Thien Lam, Jessica Kumke, Sreejana Ray, Zilia Yanira Muñoz Ramirez, Sanchita Das, Hanh Ngoc Lam

**Author notes:** Address correspondence to Hanh Ngoc Lam.

## Abstract

**2. Abstract:** *Achromobacter xylosoxidans* is an emerging pathogen characterized by high levels of antibiotic resistance and increasing infection rates worldwide. This motile, opportunistic pathogen is widely distributed in the environment and can cause various infections, including pneumonia, bacteremia, endocarditis, meningitis, and others. In this study, we analyzed the population structure, antibiotic resistance profiles, and virulence factors of over 200 publicly available genomes. Core genome analysis revealed that *A. xylosoxidans* is highly adaptable, possessing a relatively small core genome. Antibiotic susceptibility testing of isolates from the United States revealed high resistance to multiple antibiotics. Our data show that imipenem/relebactam (IMR) is as effective against *A. xylosoxidans* as imipenem (IMI) alone, indicating that relebactam does not inhibit β-lactamase activity in *Achromobacter.* The species features multiple secretion systems, including the Type III secretion system (T3SS) of the YscN family, which is similar to those found in *Bordetella pertussis* and *Pseudomonas aeruginosa*. Isolates collected from the same patients showed changes in cytotoxicity, flagella motility, biofilm and antibiotic resistance suggesting its dynamic adaptation to host environment. Intra-host evolved isolates, NIH-010, NIH-016 and NIH-018 demonstrated the loss of flagellar motility and variable cytotoxicity while exhibiting increased antibiotic resistance and enhanced biofilm formation. Sequence analysis suggests that NIH-016-3 has tyrosine to histidine mutation at position 330 near the FlhF Guanosine triphosphate (GTP)-binding domain that may affect flagellar assembly. Interestingly, virulence assays showed significant variation in the ability of different *A. xylosoxidans* isolates to induce cell death in *in vitro* models, suggesting its dynamic adaptation to host environment.

**3. Importance:** This study provides a comprehensive examination of *Achromobacter xylosoxidans*, an emerging pathogen of global concern due to its high antibiotic resistance and increasing clinical relevance. By analyzing over 200 genomes, we offer critical insights into the population structure, resistance mechanisms, and virulence factors of this species. The identification of a small core genome underscores its potential for genomic plasticity. The existence of multiple secretion systems highlights the great capacity of *A. xylosoxidans* as a pathogen. Variations in virulence among *A. xylosoxidans* isolates indicate the complexity of this pathogen underscoring the need for further studies on its virulence mechanisms. Evolution within the host includes the loss of motility-associated systems and enhanced antibiotic resistance and biofilm formation. This work showed that *A. xylosoxidans* are resistant to relebactam when combined with imipenem a combination effective in other bacteria. These findings emphasize the urgent need for targeted therapeutic strategies to combat this opportunistic pathogen.

## 4. Introduction

The genus *Achromobacter* is a non-lactose fermenting, Gram-negative, motile bacterium recognized as an emerging pathogen in cystic fibrosis (CF) patients (1). It belongs to the β-proteobacteria class. While *Achromobacter* species are ubiquitously found in water and soil, and can grow in the plant rhizosphere, *A. xylosoxidans* isolates are primarily clinically derived (2). This bacterium was first isolated in 1971 from an ear infection (3). *A. xylosoxidans* can cause a wide range of infections, including pneumonia, bacteremia, endocarditis, meningitis, and others (4, 5). These infections, which predominantly affect immunocompromised individuals often lead to severe outcomes (4). Colonization with *A. xylosoxidans* is considered a marker of disease severity in CF patients (6) and is associated with a decline in respiratory function (7) and with a greater risk of death in lung transplant recipients (8). The persistence of *A. xylosoxidans* in both extreme environments and host organisms may be attributed to its genetic diversity and adaptability, although this aspect remains poorly understood (9). Recent virulence studies have utilized various clinical isolates to explore its pathogenic mechanisms (10–12). As of the time of this study, nearly 200 genomes of *A. xylosoxidans* were available on NCBI. However, comprehensive analyses of the differences among isolates with respect to their genomic content, virulence potential, and antibiotic resistance profiles are lacking.

Clinical isolates possess unique genomic features that differentiate them from environmental isolates and contribute to virulence, such as the type III secretion system (T3SS), a polysaccharide island, adhesion-related proteins, and several putative toxins (13). In addition to these genetic elements, the Type VI Secretion System (T6SS) has been recently identified to contribute to virulence alongside the T3SS (14, 15). The T3SS enables bacteria to deliver effector proteins directly into the cytosol of animal or plant host cells (14). These systems have diversified into seven families, spreading across Gram-negative bacteria through horizontal gene transfer (16). The T6SS facilitates bacterial competition and functions as a virulence factor in various pathogenic bacteria (17, 18). In *A. xylosoxidans*, the activity of the T6SS contributes to the internalization in immune cells wherein it expresses cytotoxic effectors (11, 18). Some environmental isolates also contain phage regions encoding components of type IV secretion systems (T4SS), and several genetic islands that may enhance survival in host-free environments (19). In addition to secretion systems, motility is a major virulence factor in pathogenic bacteria with *A. xylosoxidans* producing peritrichous flagella (20). Loss of functional flagella is commonly associated with reduced virulence (21).

The genus *Achromobacter* is well-known for its inherent resistance to a broad range of antimicrobial agents (22) and its ability to acquire resistance to commonly used treatments for respiratory infections, such as fluoroquinolones, carbapenems or azithromycin (22, 23). *A. xylosoxidans* encodes at least 50 drug resistance – associated genes, including β-lactamases, efflux pumps, and transposons (24). Seventeen different efflux pumps have been identified in *A. xylosoxidans,* which may contribute to its intrinsic resistance (24). However, to date, only four of these efflux pumps have been characterized. Notably, the observed susceptibility profiles often poorly correlate with the presence of putative resistance genes (24). These findings highlight the need to investigate the prevalence and mechanisms of antibiotic resistance (AR) among *A. xylosoxidans* isolates (25–28).

In this work, we examined the genomic diversity of all available genome sequences of *Achromobacter xylosoxidans*. Phylogenetic analysis was performed on a curated set of the available genomes, totaling 178, including those from NCBI and our recently collected clinical isolates from the NIH Clinical Center. Given the critical role of secretion systems and motility in virulence, we analyzed the presence of all secretion systems (types I to IX) and flagellar system in clinical isolates. Phylogenetic analysis was performed to classify the T3SS family of *A. xylosoxidans*. To examine the heterogeneity of phenotypic traits in *A. xylosoxidans* across major virulence systems, we performed assays to assess the ability of NIH clinical isolates to induce cytotoxicity, form biofilms, and exhibit flagellar motility.

## 5. Results

### Genomic diversity of *Achromobacter*

To examine the diversity among *A. xylosoxidans* isolates, we analyzed the whole-genome sequencing data from 48 *Achromobacte*r spp. isolates (including 34 *A. xylosoxidans*, and other *Achromobacter* species collected at the NIH clinical center in the previous study (29) referred to as NIH isolates) and 197 *A. xylosoxidans* genomes from NCBI (referred to as NCBI isolates). Although our focus was on *A. xylosoxidans*, we included other *Achromobacter* species from our NIH isolates as references to assess their clustering relative to *A. xylosoxidans*. The 197 NCBI isolates were predominantly from the United States (n = 51), Germany (n = 39), Russia (n = 19), China (n = 12), and Australia (n = 11) (Figure 1). In this dataset, *A. xylosoxidans* isolates were primarily collected from respiratory sources (e.g. cystic fibrosis patients), followed by environmental samples (Figure 1A). Among the NIH isolates, the majority of samples were collected from respiratory sources, with bloodstream isolates being the second most common (Figure 1B).

**Figure 1.**
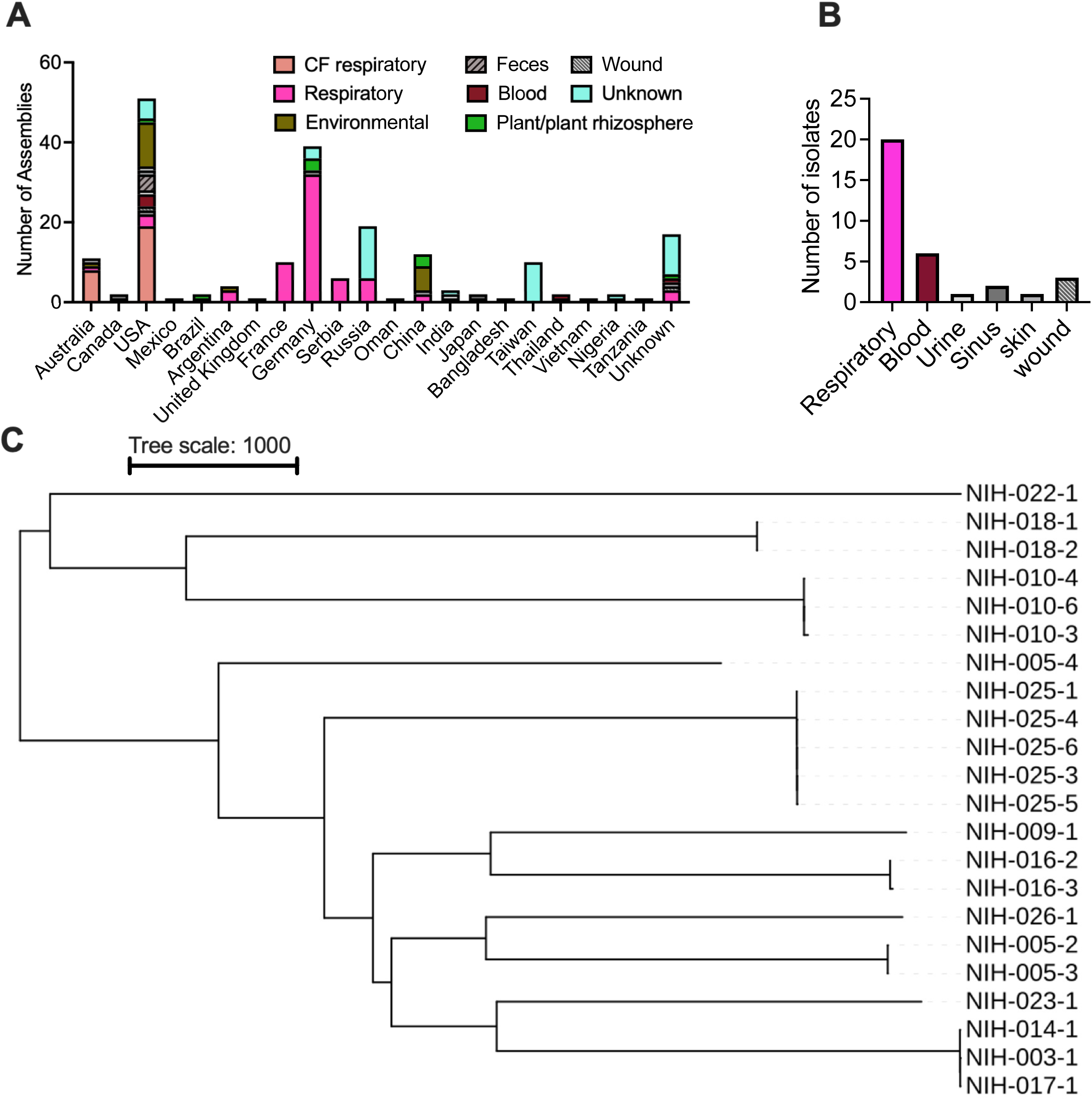
Geographic distribution, collection sources, and phylogenetic relationships of *Achromobacter xylosoxidans* isolates used in this study. (A) Number of isolates collected from different countries worldwide and their respective collection sources in the NCBI dataset. (B) Collection sources of *A. xylosoxidans* NIH isolates included in this study. Respiratory isolates include both cystic fibrosis (CF) and non-CF patient samples. (C) Phylogenetic tree of 22 curated sequences of *A. xylosoxidans* NIH isolates.

From the NIH isolate sequence, genome assembly was performed and contigs greater than 500 bps with coverage of 20 or more were retained. CheckM analysis identified 10 genomes with contamination levels exceeding 5%, which were removed from further analysis (Table S1). We note that the apparent “contamination” issue is likely due to sequencing errors. These isolates originated from single colony collections and were identified as *A. xylosoxidans* based on NCBI taxonomy analysis. Therefore, in later phenotypic analyses, we added back two isolates (NIH-016-1 and NIH-018-3) to get a complete picture of the phenotypic changes of isolates collected from the same patients. This resulted in 38 genomes from the NIH isolate dataset. For the NCBI genomes, those with more than 300 contigs were excluded, and contigs shorter than 500 bps were removed, one additional genome was further excluded from the NCBI dataset due to low genome completeness (less than 70%) resulting in a set of 140 NCBI sequences (Table S2). This yielded a final dataset of 178 genomes for downstream analysis.

Phylogenomic analysis of all 178 isolates revealed significant diversity, with isolates largely being nonclonal (Figure S1). NIH isolates were scattered among NCBI isolates, and clustering did not correlate with geographical locations or sample sources. A group of environmental isolates formed a distinct cluster (except for NIH-006-1), while others clustered with NIH clinical isolates, underscoring the complexity of *A. xylosoxidans*. Non-*A. xylosoxidans* isolates clustered together, suggesting the ability of this analysis to differentiate *Achromobacter* species. We note that two isolates classified as *A. xylosoxidans* in the previous study - NIH-019-1 and NIH-021-1 (29) – were clustered with non-*A. xylosoxidans*. A closer examination of the taxonomy analyses associated with the SRA sequences on NCBI along with TYGS analysis (30), suggests that they are unlikely to be *A. xylosoxidans*. We performed the average nucleotide identify (ANI) analysis and confirmed that the two isolates are not *A. xylosoxidans* (Figure S2). Therefore, we excluded them from subsequent analyses involving *A. xylosoxidans*. This resulted in a total of 22 *A. xylosoxidans* NIH isolates with the genome sequences. In co-ancestry analysis, there are a few cases with some level of shared ancestry from environmental samples. However, most isolates are unrelated to each other, suggesting that they are genetically diverse and likely evolve independently (Figure S3). We then examined the core genome of all 162 *A. xylosoxidans* isolates (140 NCBI and 22 NIH isolates). Using the Panaroo pipeline, we identified 2,216 genes present in at least 99% of the isolates, representing 34% of total genes in *A. xylosoxidans*, and 3,490 genes present in at least 95% of the isolates, representing 54% (Table 1). The remaining 66% of genes were classified as accessory genes, highlighting the genomic plasticity of this species.

**Table 1:**
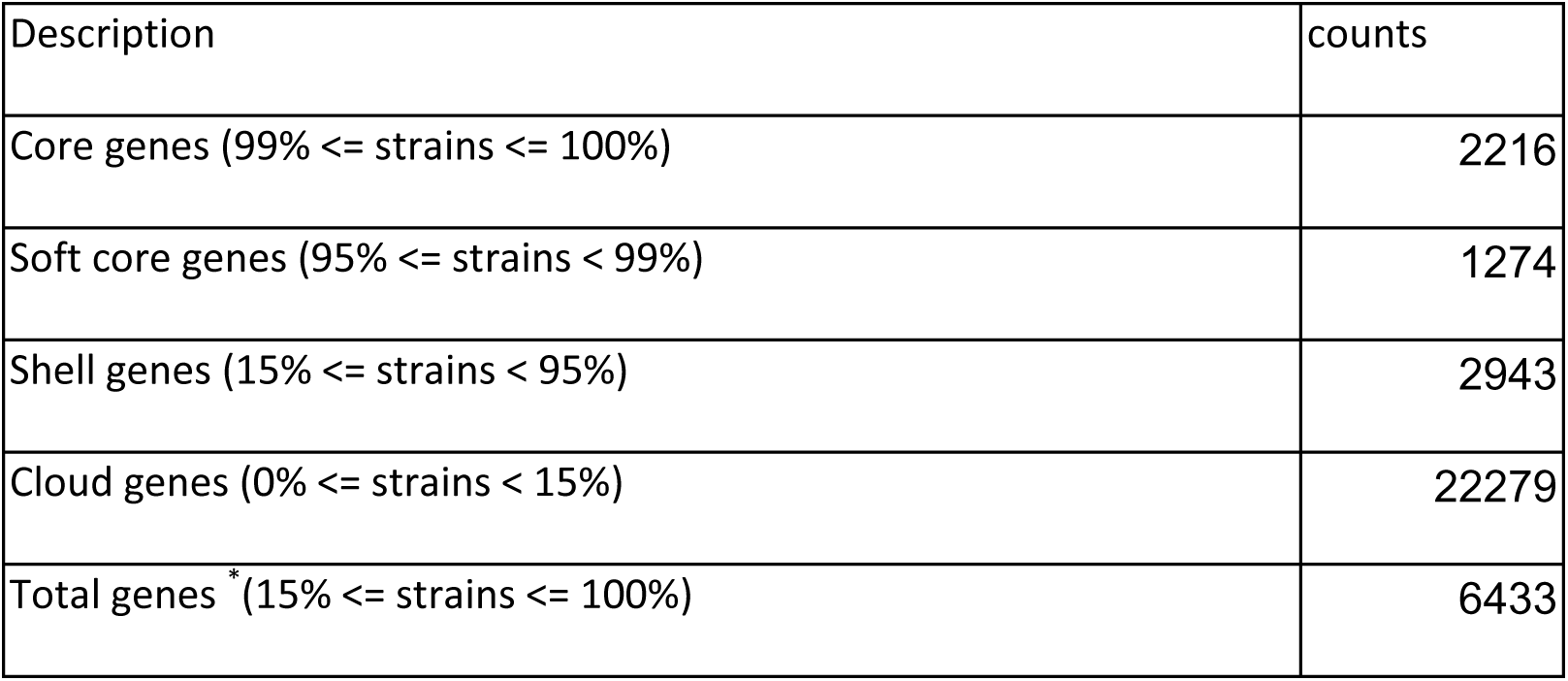
Core genes in Achromobacter xylosoxidans. * Rare genes that occurs in less than 15% of isolates are not counted in the total genes to avoid underestimate the core gene for each genome.

We also specifically looked at the phylogenetic analysis of the 22 *A. xylosoxidans* genomes from the NIH isolates (Figure 1C). In this data, isolates collected from the same patient over time have the same middle number. For example, NIH-016-2 and NIH-016-3 were from the same patients; or NIH-010-3, NIH-010-4, NIH-010-6 were from the same patients. Initial phylogenetic analysis using two methods, kSNP or Panaroo followed by RaxML (see Methods section for details) yielded identical phylogenetic trees for the NIH isolates data. Using Single Nucleotide Polymorphisms (SNPs), we built a phylogenetic tree of these isolates based on their SNP differences (Figure 1C). This tree showed the same relationships among isolates collected from the same patients as observed with the other two methods. The analysis showed that isolates collected from the same patients clustered together, indicating genetic similarity and suggesting persistence and evolution within the same host over time. The only exception was NIH-005-4, which did not cluster with NIH-005-2 and NIH-005-3. NIH-005-4 was collected from sputum in 2014, whereas NIH-005-2 and NIH-005-3 were collected from the perirectal region and bronchoalveolar lavage, respectively, in 2016.

### Antibiotic resistance in *Achromobacter*

Antibiotic susceptibility data of NIH isolates revealed a prevalence of multidrug-resistant *Achromobacter* spp. among the 12 antibiotic tested, including CZA (Ceftazidime/Avibactam), C/T (Ceftolozane/Tazobactam), IMR (Imipenem/Relebactam), MERO (Meropenem), AZT/ATM (Aztreonam), FEP (Cefepime), TAZ (Ceftazidime), CIP (Ciprofloxacin), IMI (Imipenem), LEVO (Levofloxacin), PT4/TZP (Piperacillin/Tazobactam), and SXT (Trimethoprim/Sulfamethoxazole) ((29), Figure 2A). Many AR genes were identified in the NIH isolates using the Comprehensive Antibiotic Resistance Database (CARD database); however, these genes could not fully explain the AR in certain isolates (29). Two notable isolates, NIH-016-2 and NIH-016-3, were collected from the same patient. However, NIH-016-3 was resistant to 11/12 antibiotics (all antibiotic tested except SXT), while NIH-016-2 was resistant to only 3/12. Antimicrobial Resistance analysis using CZ ID website (31) showed that NIH-016-2 and NIH-016-3 isolates carried similar sets of AR systems, including efflux pumps (AdeFGH, AxyXY-OprZ, MexAB-OprM, MuxABC–OpmB, MexGHI-OpmD, MexCD-OprJ, MexJK-OprM, MexPQ-OpmE), OXA-114h *β*-lactamase, AXC *β*-lactamase, and CatB. Thus, we re-sequenced both NIH-016-2 and NIH-016-3, as well as NIH-016-1, which has the same level of antibiotic resistance as NIH-016-3 and was collected after NIH-016-3. Using both short-read and long-read sequences, we were able to assemble complete genomes of the three isolates. Then, variant analysis was done using NIH-016-2 as a reference (Table 3). Interestingly, the variant analysis identified a T>G single nucleotide polymorphism (SNP) at position 29 of *axyZ*, resulting in a nonsynonymous mutation (V29G) in both NIH-016-3 and NIH-016-1. AxyZ is a negative regulator of AxyXY-OprZ RND efflux pump. This mutation (V29G) is associated with overexpression of AxyXY-OprZ efflux pump, which may be involved in resistance against multiple antibiotics including tobramycin, gentamicin, cefepime, ofloxacin, and levofloxacin (32). SNPs that may be involved in antibiotic resistance were identified in NIH-016-03 and NIH-016-01, and among these, three were of particular interest (Table 3). A T>A SNP at position 695 of *mutL* resulted in a non-synonymous mutation (L232Q). MutL and MutS are components of the mismatch repair (MMR) system, and mutations in these genes, especially MutL, are associated with a hypermutator phenotype and increased antibiotic resistance in *P. aeruginosa* (33). Another variant, T>C at position 2228 in the *mfd* gene (ACP6EQ_07680) caused an L743P substitution. *mfd* has been linked to elevated mutagenesis rate and the development of antibiotic resistance in several bacterial species (34). Finally, a C>T SNP at position 592 of a gene encoding an ABC transporter substrate-binding protein (ACP6EQ_16040) produced an R198C. ABC transporters play important roles in regulating the uptake and efflux of various substrates, including antibiotics (35). Overall, NIH-016-3 and NIH-016-1 appear to be hypermutator strains that allow the accumulation of additional mutations, thereby enhancing the antibiotic resistance of these isolates.

**Figure 2:**
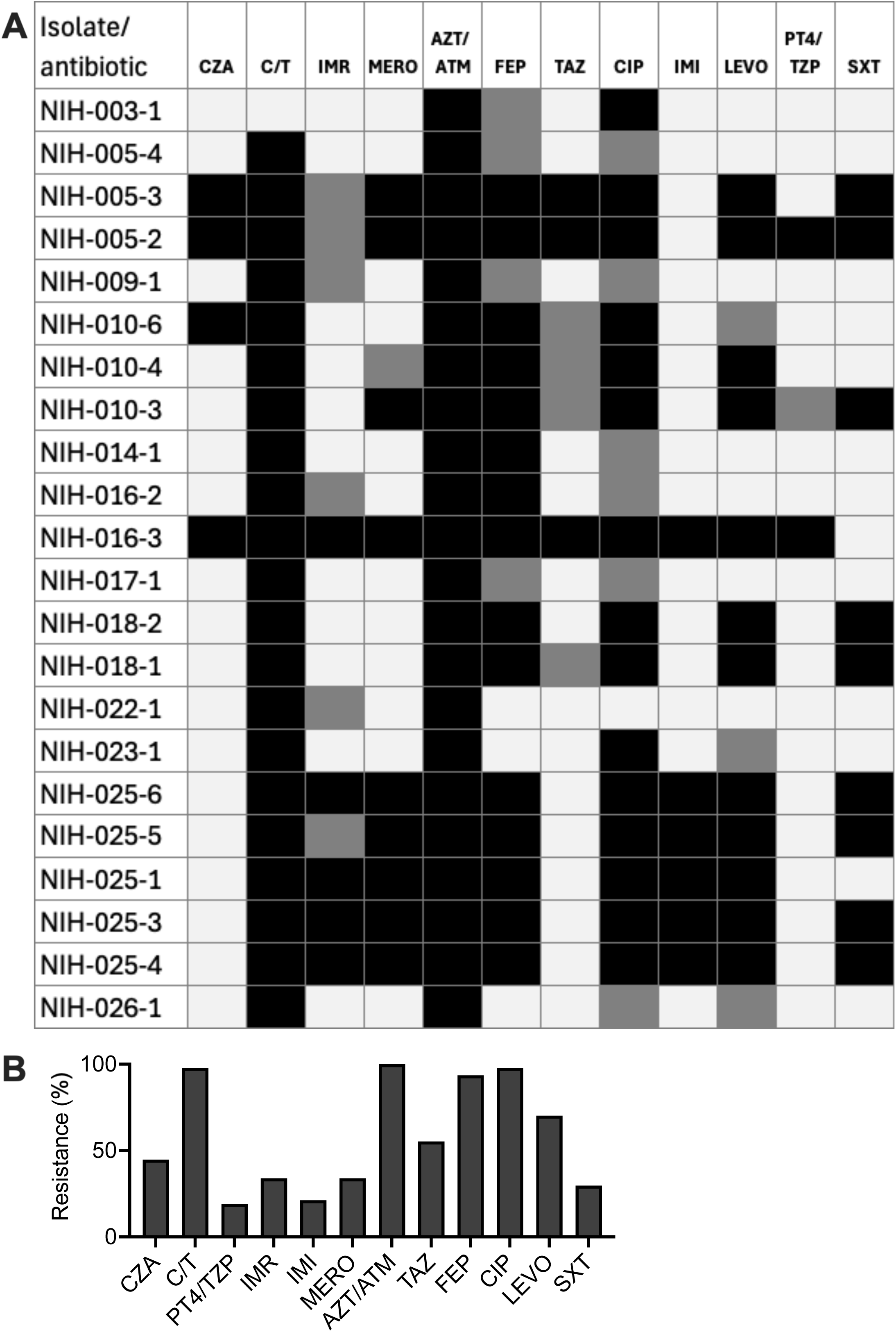
Frequency of antibiotic resistance in NIH isolates. (A) Antibiotic resistance in NIH clinical isolates: CZA (Ceftazidime/Avibactam), C/T (Ceftolozane/Tazobactam), IMR (Imipenem/Relebactam), MERO (Meropenem), AZT/ATM (Aztreonam), FEP (Cefepime), TAZ (Ceftazidime), CIP (Ciprofloxacin), IMI (Imipenem), LEVO (Levofloxacin), PT4/TZP (Piperacillin/Tazobactam), and SXT (Trimethoprim/Sulfamethoxazole). Black: resistance, gray: intermediate, white: susceptible. (B) Percentage of isolates resistance to each antibiotic was shown.

### Secretion systems in *Achromobacter xylosoxidans* clinical isolates

Previous studies have shown that *A. xylosoxidans* can express multiple virulence factors, including secretion systems and flagella (14, 23). We performed a genome scan for secretion systems and appendages using MacSyFinder to identify secretion systems and flagella in the 22 *A. xylosoxidans* NIH isolates (Table 2). The reference genome (FDAARGOS_1091) and two additional clinical isolates (GN050 and GN008) were included for comparison (Table 2). All isolates possessed T1SS, T2SS, T5SS, and bacterial Tight adherence Secretion System (TadSS). T6SS was present in all isolates, and T4SS was present in all isolates except NIH-009-1. Interestingly, although a number of isolates were collected from the same patients over time, most of them still carry key flagella and T3SS genes. Only one isolate, NIH-023-1 lost both flagellar and T3SS genes.

**Table 2:**
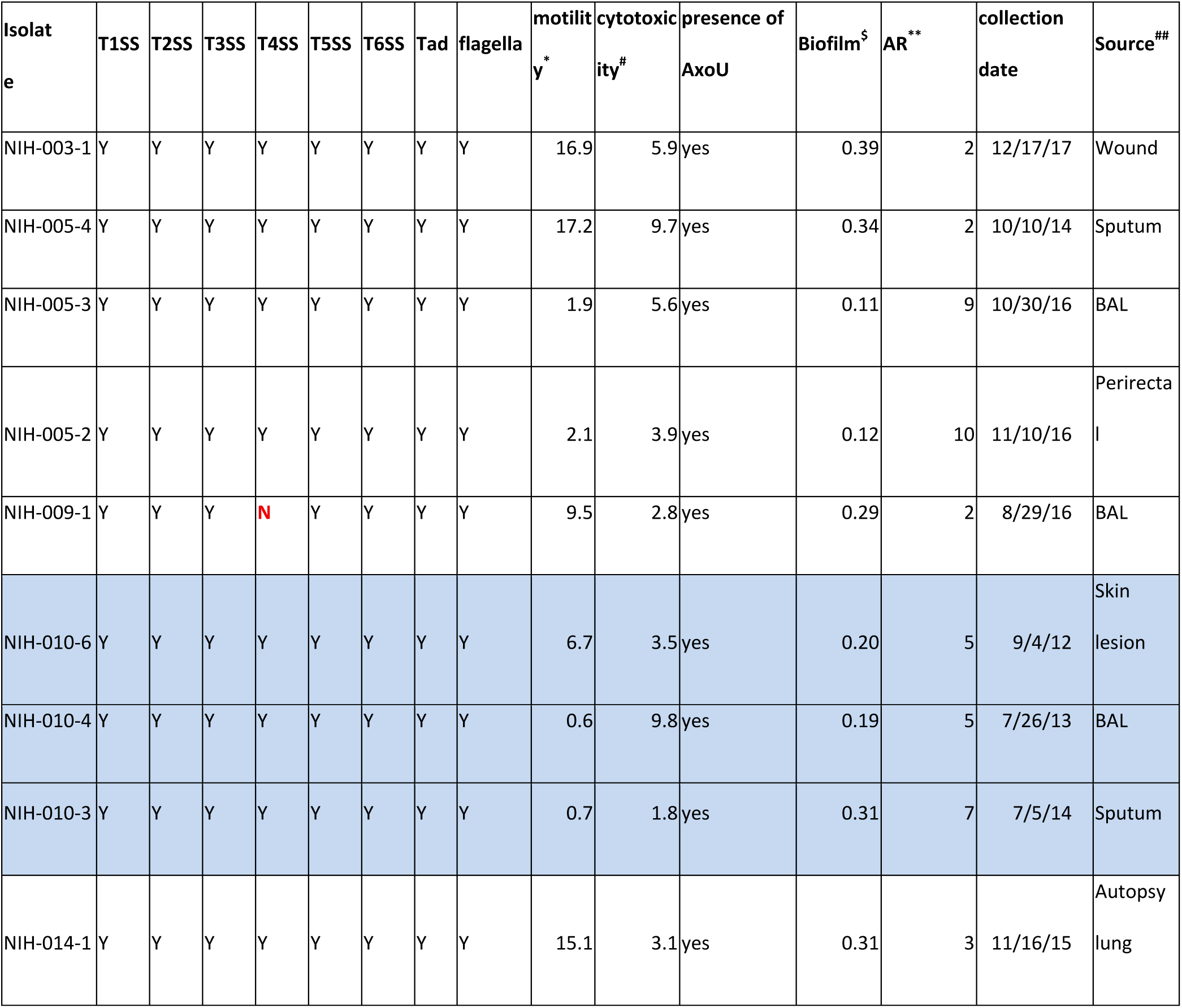
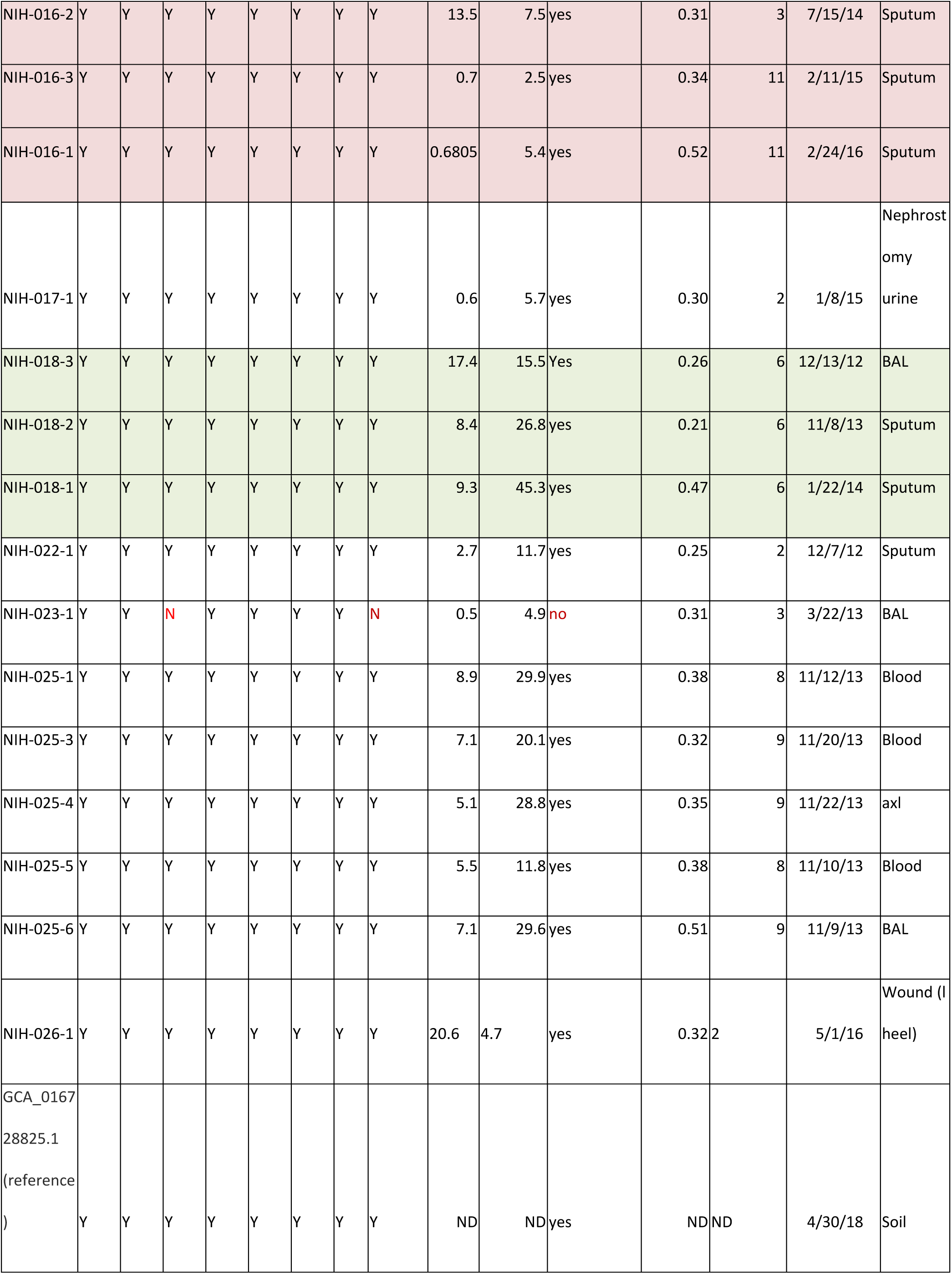
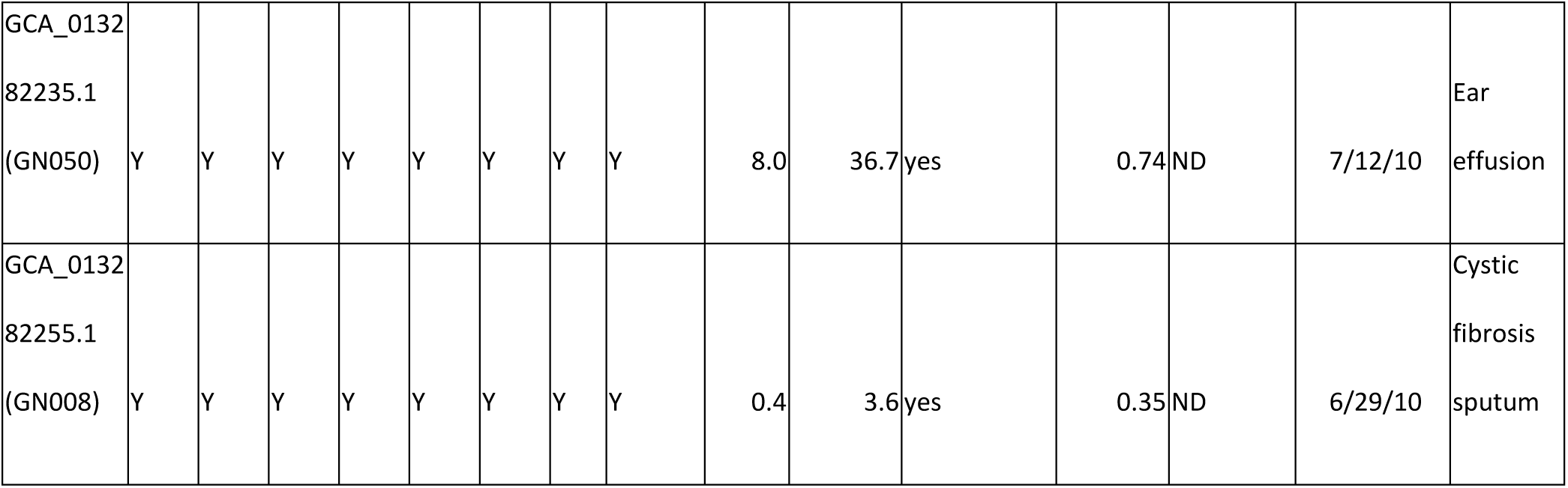
Summary of secretion system, motility, cytotoxicity, and biofilm of *A. xylosoxidans* isolates. Three intra-host isolate series were highlighted in different colors. NIH-016-1 and NIH-018-3 were included in the table for the analysis of intra-host samples, while the genome sequences were omitted in the phylogenetic analysis due to contamination issue. *motility in mm^2^ ^#^cytotoxicity: percent cell death at 5 hours after infection ^$^ biofilm: Absorbance 595nm of crystal violet stained biofilm **AR: Total number of antibiotics that each isolate is resistant to out of 12 tested ^##^ BAL: Bronchoalveolar lavage, AXL: Autopsy of the lung

**Table 3:**
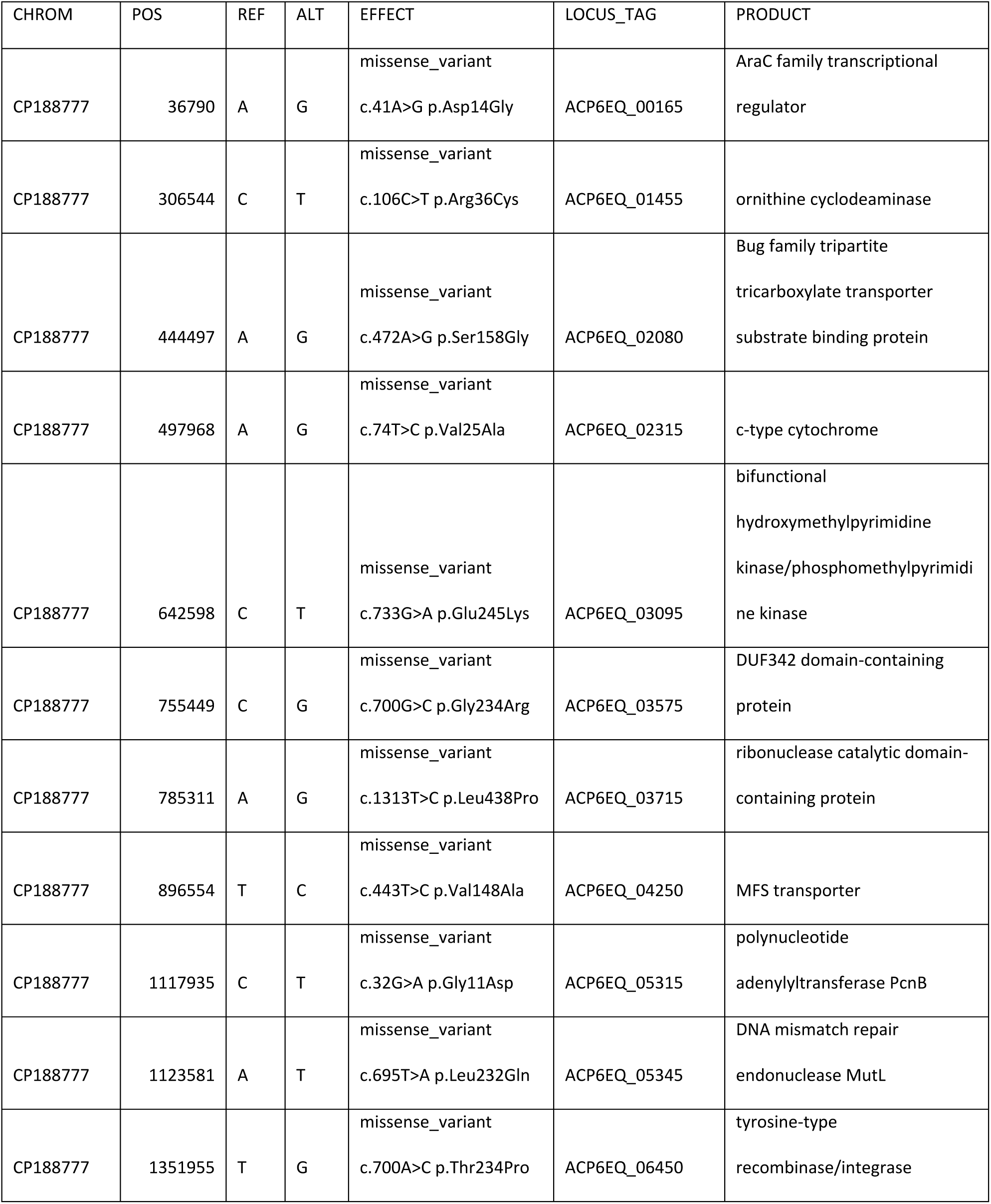
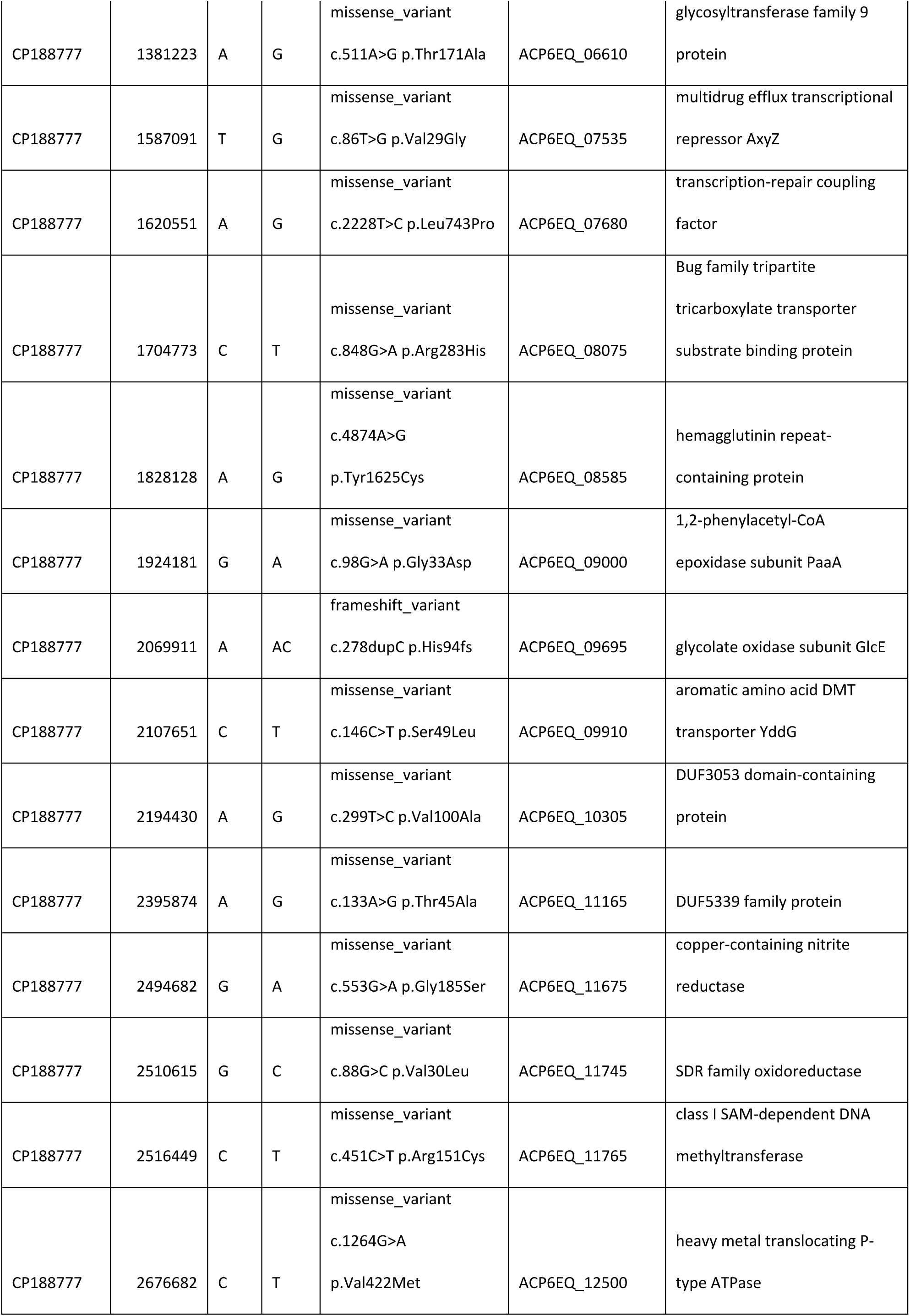
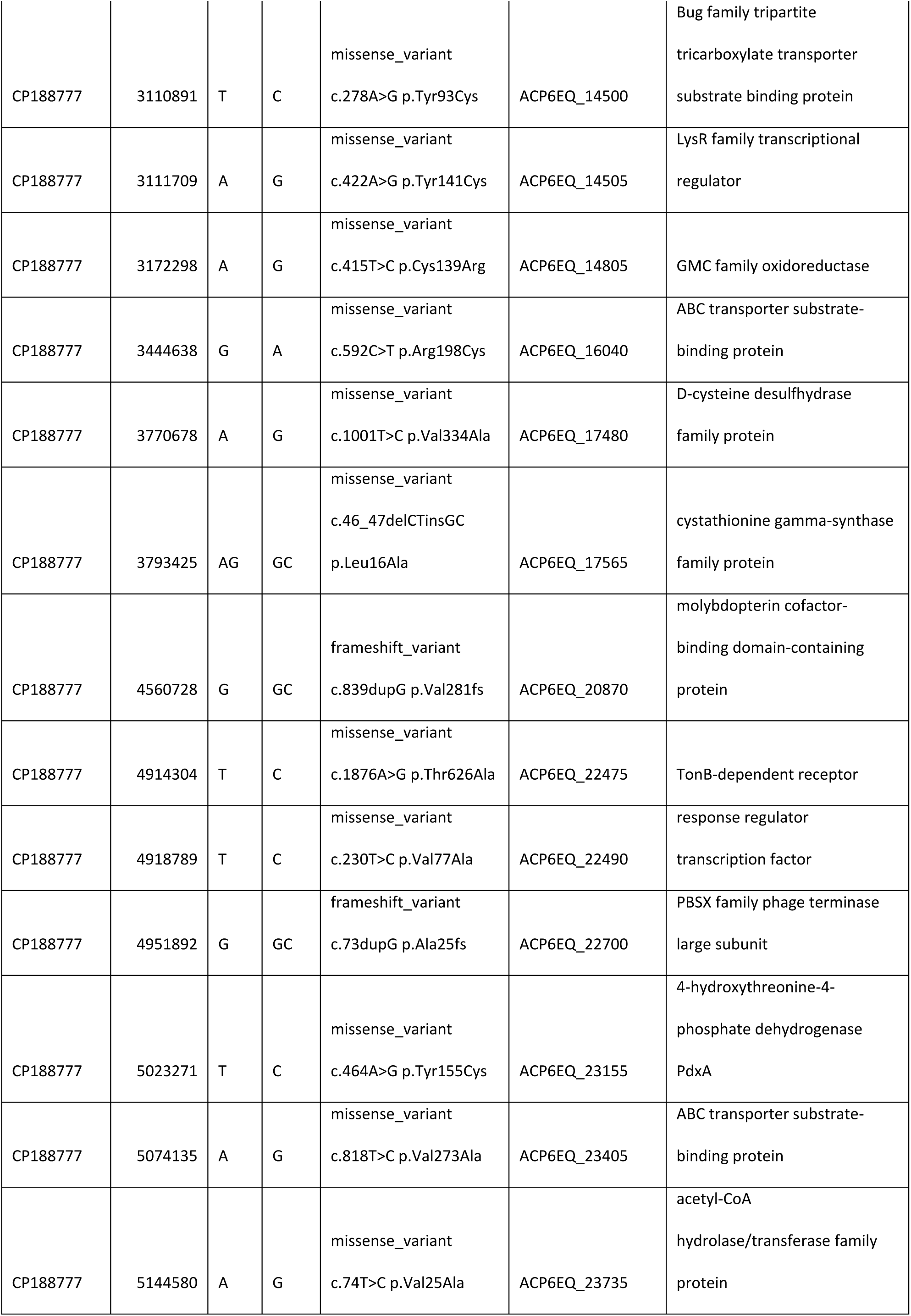
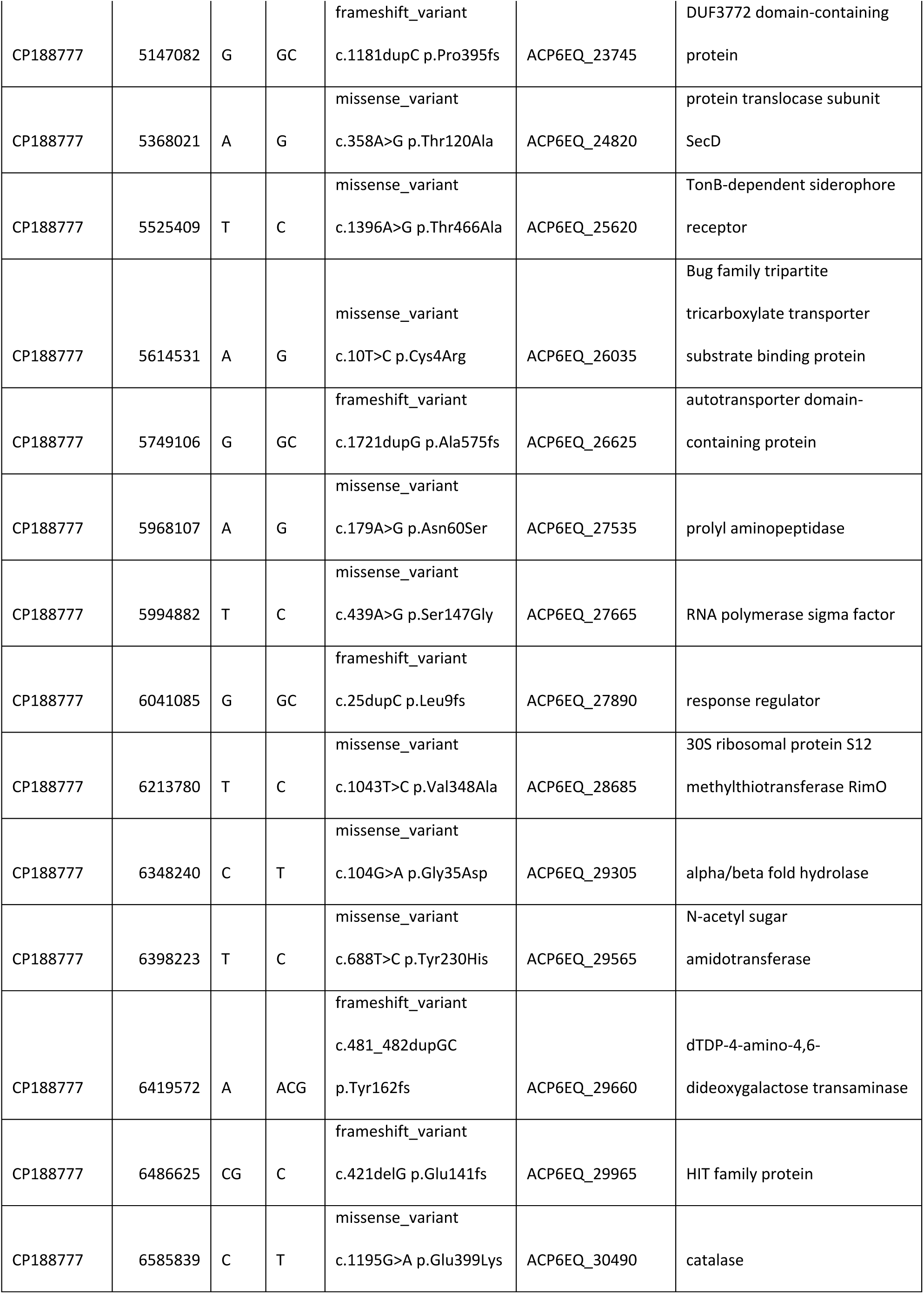
Changes of the genome between NIH-016-2 and NIH-016-1, NIH-016-3. NIH-016-2 was used as a reference. The mutations listed are the common mutations between NIH-016-3 and NIH-016-1, compared to NIH-016-2.

The type III secretion system is an important virulence mechanism contributing to cytotoxicity in *A. xylosoxidans* (11, 14). Phylogenetic analysis showed that the T3SS in *A. xylosoxidans* belongs to the Ysc T3SS family (Figure 3A), which includes T3SS from other pathogens such as *P. aeruginosa*, *Yersinia pseudotuberculosis,* and *Bordetella pertussis*. Structural modeling of the APTase YscN from *A. xylosoxidans* showed high similarity to that of *P. aeruginosa* PA14, with a root-mean-square deviation (RMSD) of 0.488 A° (Figure 3B, C). This structural similarity further supports the classification of *A. xylosoxidans* T3SS within the Ysc family.

**Figure 3:**
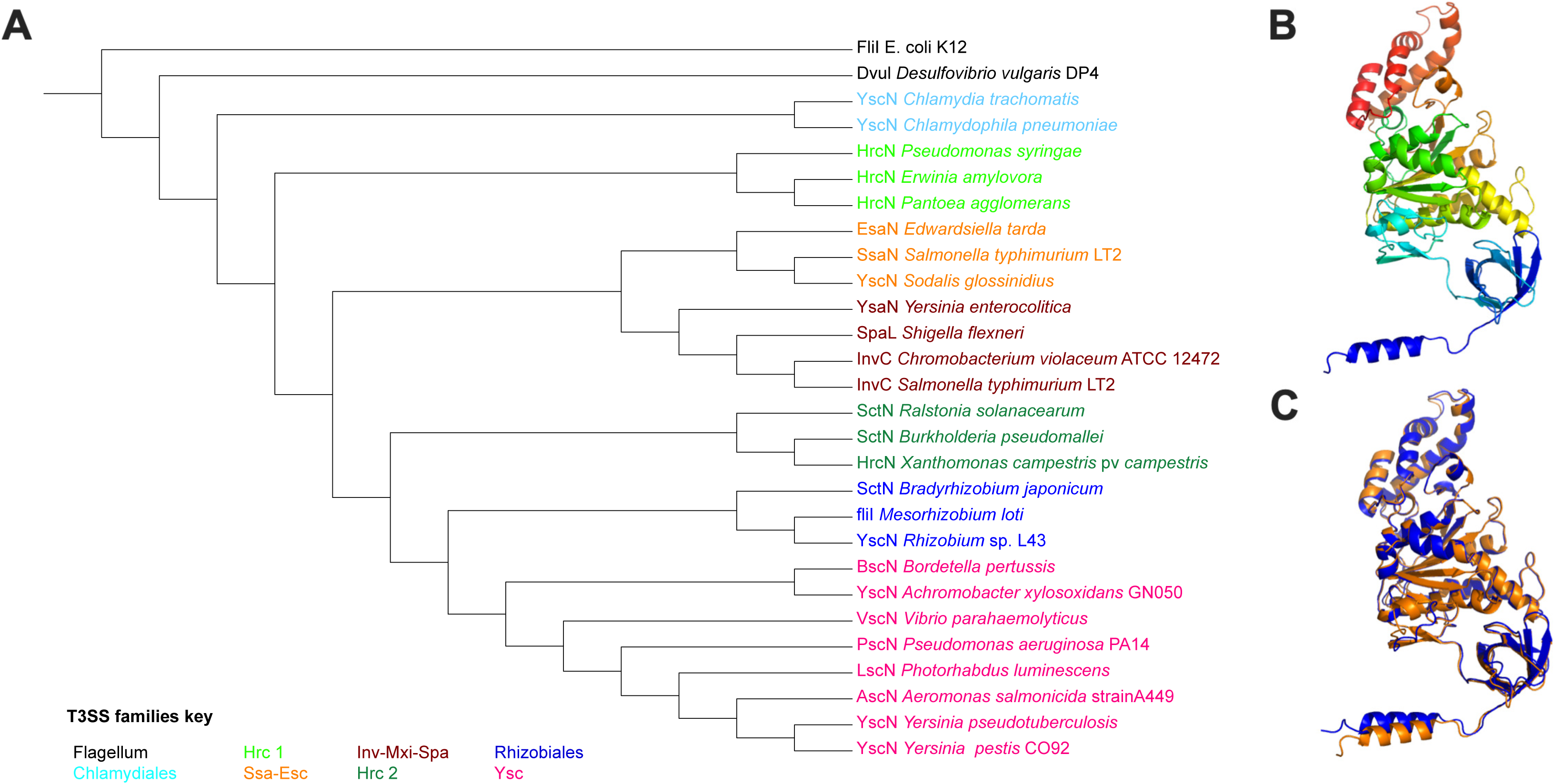
Type III secretion system (T3SS) phylogeny and structure. (A) Phylogenetic tree of T3SS families constructed from aligned amino acid sequences of SctN and its homologs. (B) Predicted structure of YscN from *Achromobacter xylosoxidans*. (C) Superimposed predicted structures of *A. xylosoxidans* YscN (orange) and *Pseudomonas aeruginosa* PA14 PscN (blue).

### Cytotoxicity in *Achromobacter*

To assess the cytotoxicity of the NIH isolates, we infected J774a.1 macrophages with each isolate at an MOI of 7 and measured cell death over time. We included two clinical isolates, GN050 and GN008, which were previously reported to exhibit differential cytotoxicity, as controls (14). The result showed that the 24 isolates have variable cytotoxicity levels with NIH-018-1 being the most cytotoxic, exceeding that of GN050 (Figure 4 A,B). Among clinical isolates no measurable difference was detected between the isolate source. Among the isolates collected from the same patients, NIH-010 and NIH-016 had minimal cytotoxicity, while all members of the NIH-025 series are cytotoxic. In the NIH-018 series, the bacterial isolates appeared to have increased cytotoxicity over time, as isolates collected later in the infection were more severe.

**Figure 4:**
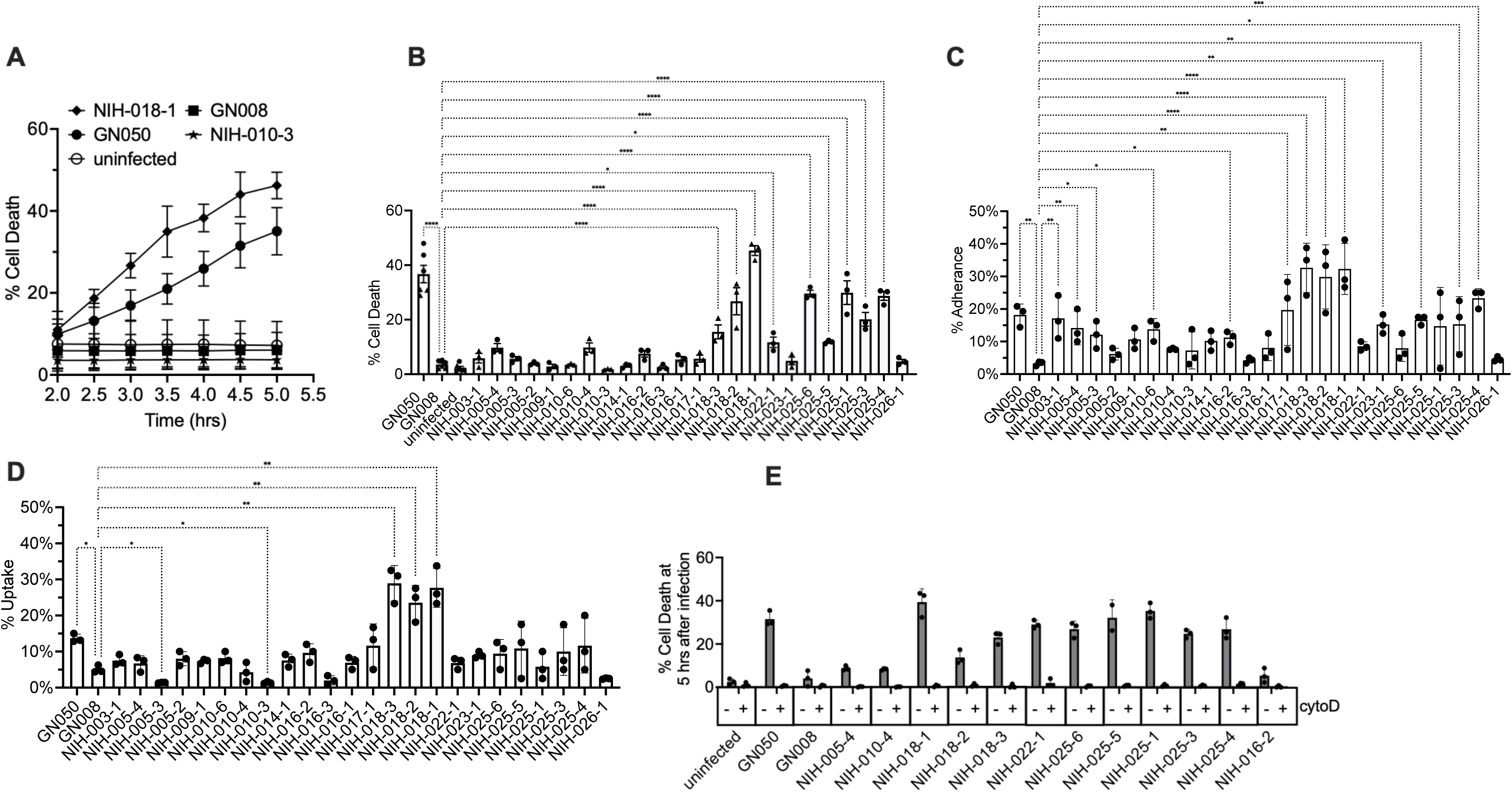
Cytotoxicity, adherence and uptake of *A. xylosoxidans* clinical isolates in macrophage. (A) Cell death of J774A.1 macrophages infected with selected *A. xylosoxidans* clinical isolates over time. (B) Cell death of J774A.1 macrophages infected with different *A. xylosoxidans* clinical isolates at 5 hours after infection (hpi). (C) Adherence and (D) internalization of selected *A. xylosoxidans* isolates. Percent of inoculum is shown. (E) Cell death of J774A.1 macrophages infected with cytotoxic isolates in the presence or absence of cytochalasin D. Percent cell death, calculated as the ratio of dead to total cells, is shown. Different symbol shapes represent independent biological replicates. One-way ANOVA with multiple comparison test was used. **** P-val <0.0001, *** P-val <0.0002, ** P-val <0.002, *P-val <0.05.

To further understand the basis of cytotoxicity, we examined the adherence and uptake of *A. xylosoxidans* in macrophages. For the adherence assay, infected J774a.1 cells were washed, lysed and plated for colony forming unit (CFU) counts (Figure 4C). For the uptake assay, the experiment was performed similarly to the adherence assay, with an additional step of adding polymyxin B after one hour of infection (Figure 4 D). Overall, the most cytotoxic isolates are the those that adhered most efficiently. The NIH-018 series demonstrated the highest levels of adhesion and internalization rate among our isolates, agreeing with its overall high cytotoxicity relative to all other strains. In contrast, several of the least cytotoxic strains (e.g., NIH-05-2, NIH-010-4, NIH-010-3, NIH-014-1, NIH-016-3, NIH-016-1, NIH-022-1 and NIH-026-1) exhibited consistently low adherence and internalization in J774a.1 cells. To further assess whether cytotoxicity depend directly on bacterial uptake, we performed the cell death assay in the presence of cytochalasin D on cytotoxic isolates (Figure 4E). Indeed, cytochalasin D completely abolished the cell death induced by these isolates. Together, these results demonstrate that internalization is a key determinant of *A. xylosoxidans* cytotoxicity across clinical isolates.

### Biofilm

The capacity of a bacterium to form biofilms is a strong indicator of virulence and can be tied to the expression of other virulence factors (36–38). Additionally, biofilm formation improves the resistance of the organism to antibiotic and environmental pressures (39). Static biofilms serve as an effective method for measuring the overall capacity of a bacteria to produce biofilms and have been utilized in *A. xylosoxidans* previously (40). We therefore set out to examine biofilm formation in static culture of *A xylosoxidans* to characterize the differences between clinical isolates. *P. aeruginosa* PA01 Δ*wspF*Δ*pelA*Δ*pslBCD* was used as negative control and PA01 Δ*wspF* as a positive control (41, 42) for biofilm formation. GN050 showed robust biofilm formation compared to the control (Figure 5). Most NIH isolates demonstrated a low level of biofilm production. Only NIH-016-1, and NIH-025s displayed substantial biofilm forming capabilities over the negative control. These isolates were from sputum (NIH-016-1), blood or lung (NIH-025s), suggesting that the source of isolation does not determine whether the bacteria can form biofilm. The intra-host series, NIH-010, NIH-016 and NIH-018 do show a trend of increasing biofilm production although it’s not statistically significant from the *P. aeruginosa* negative control.

**Figure 5:**
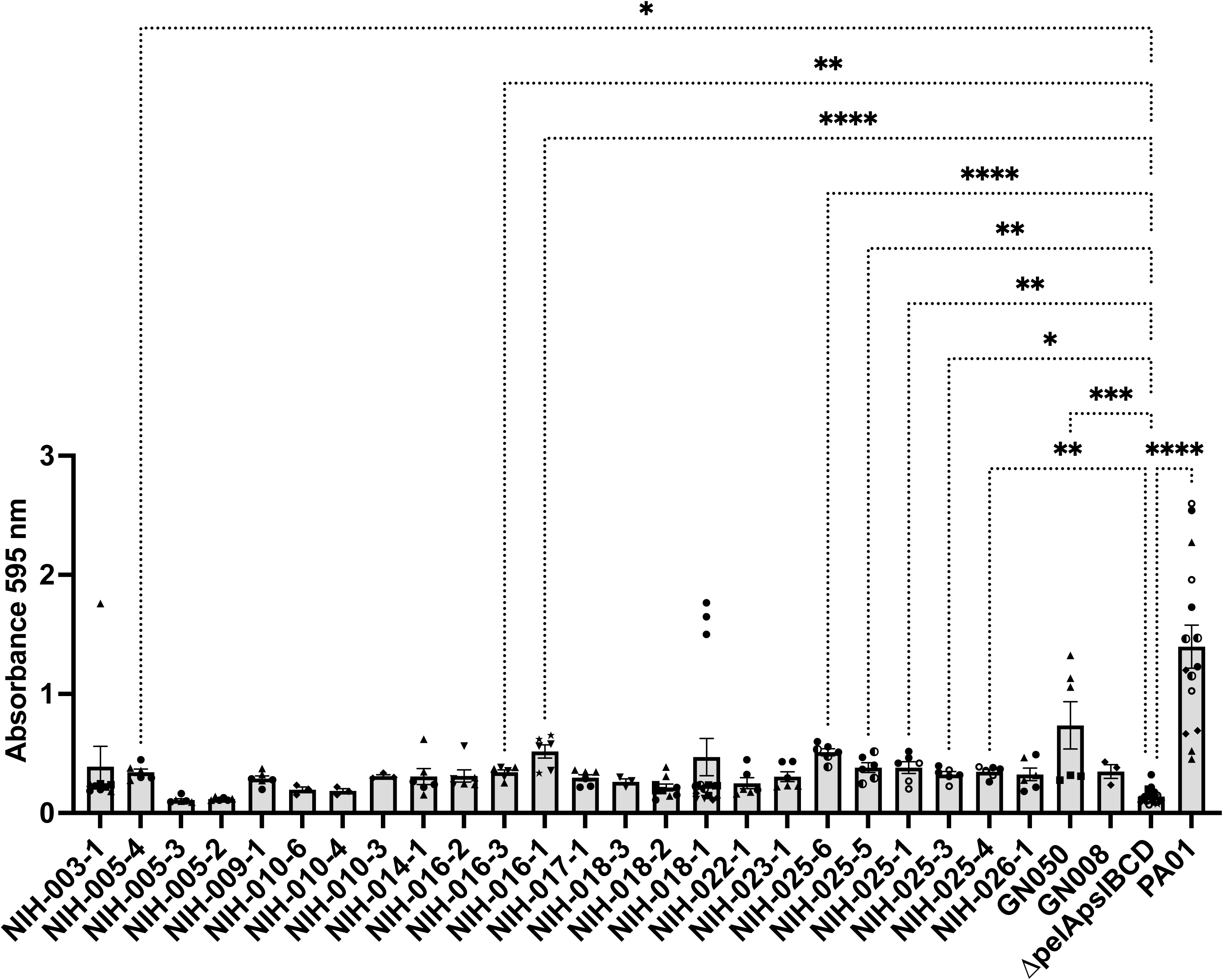
Biofilm formation by *A. xylosoxidans* clinical isolates. PA01 Δ*wspF*Δ*pelA*Δ*pslBCD* was used as a negative control and PA01 Δ*wspF* was used as a positive control. Different symbol shapes represent independent biological replicates. The experiment was done in at least three technical replicates. Kruskal-Wallis test was done followed by Dunn’s multiple comparisons to correct for multiple comparison. **** P-val <0.0001, *** P-val <0.0002, ** P-val <0.002, *P-val <0.05.

### *Achromobacter* Flagellar Motility

Flagellar motility is an important virulence factor which bacteria employ to move to the site of infection and is often coordinated with the expression of other virulence genes (43–45). During prolonged infection, mutation in flagellar genes can accumulate resulting in an overall loss of flagellar production (46). We decided to investigate the flagellar motility of our *A. xylosoxidans* clinical isolates to determine if our strains demonstrate any changes in motility. Our results demonstrated substantial differences in flagellar motility between strains even from isolates collected from the same patients (Figure 5A). The most motile isolates are NIH-003-1, NIH-005-4, NIH-014-1, NIH-016-2, NIH-018-3, and NIH-026-1. Among all 14 respiratory isolates (NIH-005-3, NIH005-4, NIH009-1, NIH010-3, NIH-010-4, NIH-014-1, NIH016-2, NIH016-3, NIH018-1, NIH018-2, NIH022-1, NIH023-1, and NIH025-6) 7 were identified as non-motile representing 53.8% of the group. Additionally, among all non-motile isolates, 80% were isolated from a respiratory source. Comparing strains isolated at different times from the same patient, NIH-005, NIH-010, NIH-016, and NIH-018 series, most demonstrated a loss of motility whereas NIH-018 series showed reduced motility. In NIH-005 series, isolates 2 and 3 were collected in the same year, two years after isolate 4. The isolates 2 and 3 don’t move while isolate 4 demonstrates high motility among all isolates. In isolates NIH-010 series, isolate 6 was more motile than isolates 3 and 4, which were isolated after isolate 6. In series NIH-018, NIH-018-3 is one of the best swimmers. The motility reduced about 50% after one and two years in the subsequent isolates, but they are still motile. These observations indicate a preference for a reduction of flagellar function in the respiratory environment.

The NIH-016 series demonstrated some of the most substantial differences in motility between isolates (Figure 6A). In order to determine what factors may be contributing to this, we investigated the genomic differences present among genomes of NIH-016 series. Multiple sequence alignment between known flagellar genes in NIH-016-2 compared to the genome of NIH-016-3 showed near perfect identity between the isolates except for two identified point mutations in *fliO* and *flhF* (supplemental table 1). The point mutation in *fliO* when translated results in a synonymous mutation. However, a T>G single nucleotide polymorphism was identified in *flhF* which resulted in a Y330H mutation FlhF (Figure 6B). This mutation also exists in NIH-016-1 which was collected after NIH-016-3. FlhF is a GTPase known to be important for the initial insertion of flagellar components in the membrane of the bacteria. Deletion or mutations of key regions of this protein are known to result in non-flagellated bacteria and thus a loss of motility (47, 48).

**Figure 6:**
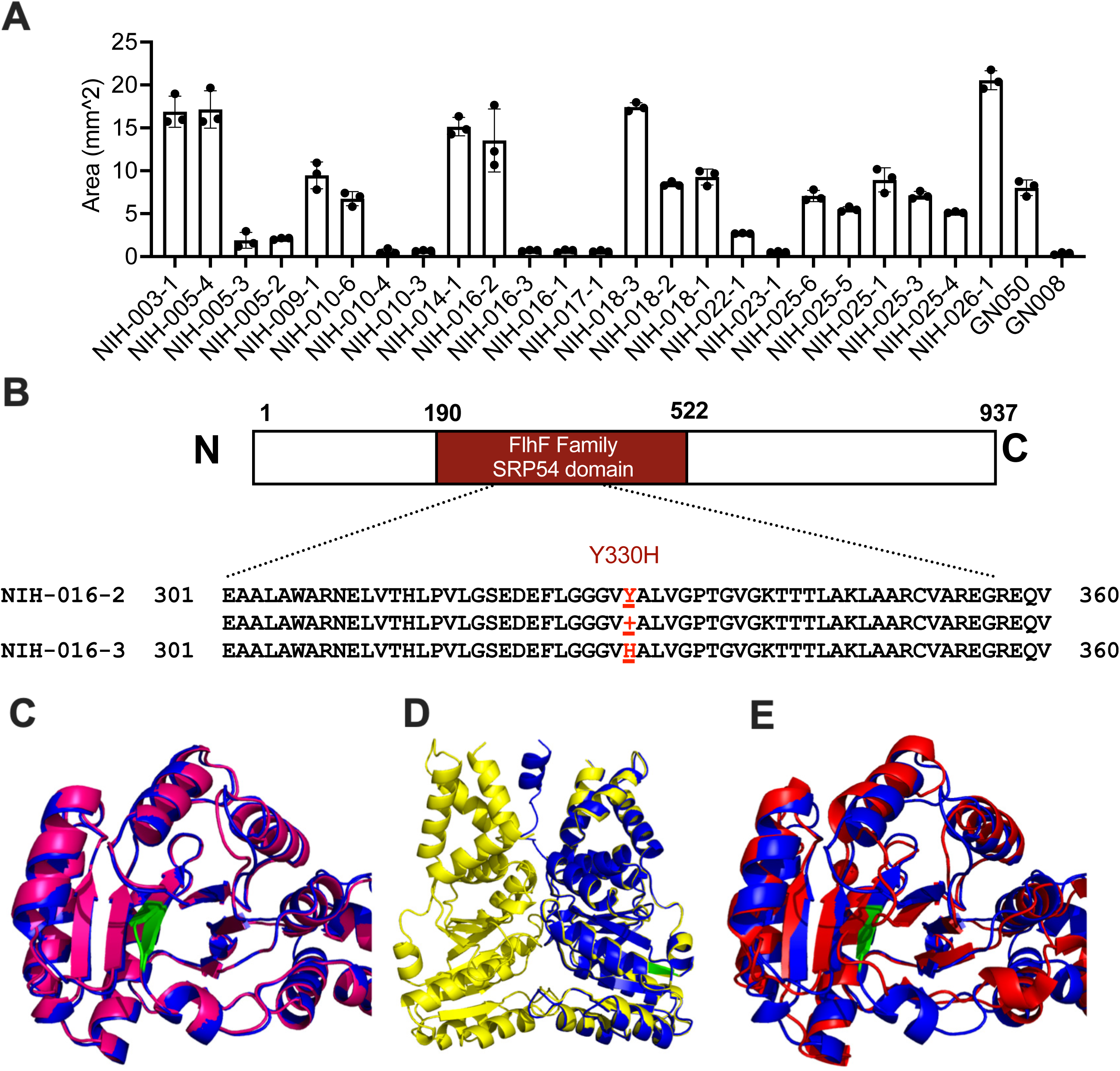
*A. xylosoxidans* motility. (A) *A. xylosoxidans* demonstrates substantial differences in flagellar motility among clinical isolates. (B) FlhF is a 937 amino acid (aa) protein that contains a FlhF family domain (aa 190-415) and an SRP54 domain (aa 327-522), based on InterProscan (109). In isolate NIH-016-3, FlhF carries a point mutation at position 330, resulting in a Y330H substitution within the SRP54 domain. (C) Mutation Y330H results in a predicted displacement of V229 (Green) as determined by modeling in Alphafold (Blue: NIH-016-2 FlhF, Red: NIH-016-3 FlhF). (D) Alignment of monomeric NIH-016-2 FlhF (Blue) to homodimerized NIH-016-2 FlhF (Yellow). (E) Alignment of NIH-016-2 (Blue) to *B. subtilis* FlhF (Pink) with bound GTP (Red).

Characterization of the protein for domains by InterProScan (49) revealed a predicted SRP54-type protein, GTPase domain which spans from 329 to 520. SRP54-type proteins represent signal recognition particles which bind GTP and are important for trafficking to the plasma membrane (48, 50). This observation is in line with the predicted function of FlhF as a GTP binding protein and its function in nucleating flagellar assembly in other bacteria (51, 52). Prediction of the FlhF protein overall via modeling in alphafold (53) revealed four distinct structures. A structure ranging from 268 to 520 comprised a GTPase domain. Alphafold modeling of NIH-016-2 FlhF and NIH-016-3 FlhF and subsequent alignment revealed a structural abnormality in NIH16-3’s FlhF which resulted in a predicted displacement of a valine at position 329 (Figure 6C). FlhF is known to homodimerize following binding to GTP (50) and therefore we generated an additional model to visualize plausible homodimer structures of NIH-0016-2 FlhF. Y330 was observed on the outside of the predicted model, indicating that the mutation may not be causing any structural disruptions directly responsible for dimerization (Figure 6D). An alternative hypothesis is the mutation may be disrupting the GTP binding domain. An alignment of the predicted *Achromobacter* FlhF structure to the known crystal structure of FlhF in *B. subtilis* showed a R^2^ of 2.082, demonstrating they are structurally similar. The known location of the GTP binding domain of *B. subtilis* FlhF is in close proximity to the β-sheet containing the Y330H mutation (Figure 6E). Using observed differences in flagellar motility we were able to identify a mutation in NIH-016-3 and NIH-016-1 which may be contributing to a loss of flagellar motility possibly by the disruption of GTP binding.

## 6. Discussion

*A. xylosoxidans* is an emerging pathogen that has garnered increasing attention in recent years. Accurate identification of *Achromobacter* species has been challenging due to limited knowledge, reliance on traditional and labor-intensive phenotypic identification methods, and the lack of comprehensive spectra from representative species within MALDI-TOF databases (54). Advances in sequencing technology and the decreasing cost of sequencing have made whole genome sequencing a feasible approach for identifying *A. xylosoxidans*. Mining these whole genome sequencing data can provide insights into the genetic diversity, antibiotic resistance mechanisms, and virulence potential of these isolates. In this study, we analyzed publicly available sequences from NCBI and our data from isolates collected at the NIH Clinical Center representing various genetic backgrounds.

Our core genome analysis revealed that *A. xylosoxidans* possesses a relatively small core genome (34–54% of its total genes when defined as being present in 99-95% of the genomes). This is smaller than the core genomes of other opportunistic pathogens such as *P. aeruginosa* and *Helicobacter pylori*. For instance, *P. aeruginosa* has approximately 40-62% core genes - 2,503 core genes (present in 100% of genomes) out of an average of 6175 genes (55), or 3,769 core genes (present in 98% of genomes) out of 6103 genes (56). Similarly, *Helicobacter pylori* has 1,227 core genes present in at least 95% of the genomes (57) out of ~1,500 total genes (58), making up 81% of its genome. The core genome phylogenetic tree of *A. xylosoxidans* appears to be non-clonal and isolates from the same geographic locations or sample sources did not cluster together. The lack of population structure and co-ancestry suggests that *A. xylosoxidans* isolates are likely evolving independently, possibly under selective pressures in different environments or hosts. Such variations may arise from diverse soil or water environments (where *Achromobacter* are usually found), or from differences in antibiotics treatment regimens, underlying host diseases, or other health conditions. It is also worth noting that we do observe an exception of a few environmental isolates that do show some level of shared ancestry. Most isolates collected from the same patients clustered together and shared co-ancestry (Figures 1C and S3) suggesting that they evolved from the same infection. However, an exception was observed with NIH-005-4, which did not cluster with NIH-005-3 and NIH-005-2, indicating that NIH-005-2 and NIH-005-3 likely represent a case of reinfection rather than a persistent infection.

The multidrug resistance of *A. xylosoxidans* is common and concerning (Figure 2). This bacterium encodes several efflux pumps, although a few have been validated. The resistance-nodulation-cell division (RND)-type multidrug efflux pump, AxyABM, showed some level of resistant to cephalosporins (except cefepime), aztreonam, nalidicic acid, fluoroquinolones and chloramphenicol (59). Recently, a novel RND-type multidrug efflux pump, AxyXY-OprZ, was identified, which expels antibiotics such as cefepime, fluoroquinolones, tetracycline or aminoglycosides including kanamycin, gentamicin, and tobramycin (60). Additionally, a recent study discovered the presence of another narrow-spectrum RND-type multidrug efflux pump, AxyEF-OprN, which pumps out fluoroquinolones, tetracyclines, and carbapenems (40). An additional study identified AxySUV in some of the *A. xylosoxidans* genomes that export fluoroquinolones (ciprofloxacin, levofloxacin), carbapenems (doripenem), tetracycline (doxycycline, minocycline), and chloramphenicol (61). In addition to the presence of various efflux pumps, intrahost evolution may contribute to increasing antibiotic resistance over time. One interesting case is the NIH-016 series. NIH-016-3, collected seven months after NIH-016-2 from the same patient, exhibits higher AR and enhanced biofilm formation compared to NIH-016-2. Phylogenetic analysis confirms their close relationship, suggesting that NIH-016-3 evolved from NIH-016-2. While both isolates carry similar sets of resistance genes identified from the CARD database and CZ ID website, NIH-016-3 demonstrates greater resistance to antibiotics. Comparing single nucleotide polymorphism of all three isolates in the series suggest that the later collected isolates accumulated mutations DNA mismatch repair systems and in several efflux pump. Hypermutators may exhibit greater resistance because their elevated mutation rate leads to additional mutations in genes that contribute to antibiotic resistance. It is possible that the hypermutator emerged first, followed by mutations in efflux pumps and transporters, ultimately resulting in broad resistance to multiple antibiotics.

In our NIH isolates, 21% of isolates are resistant to IMI while 34% isolates are resistant to IMR. Relebactam, a wide-spectrum β − lactamase inhibitor against class A (extended-spectrum β−lactamases KPC) and class C (AmpC) β−lactamases, does not seem to work in *Achromobacter*. In fact, all NIH isolates that were resistant to IMI were also resistant to IMR. In other bacteria such as *Pseudomonas* and *Klebsiella*, IMR was more effective than IMI (62, 63). This could be because *A. xylosoxidans* possesses other β−lactamases classes. NIH-016s possess AXC family carbapenem-hydrolyzing class A β−lactamase and OXA-114 family class D β−lactamase. The OXA-114 may help *Achromobacter* to escape relebactam when used in combination with imipenem. These findings highlight that *A. xylosoxidans* employ multiple AR genes and strategies, which require further research.

As secretion systems are key virulent factors in bacteria, we investigated the presence of all possible secretion systems in the NIH isolates. T1SS, T2SS, T5SS, and bacterial Tight adherence Secretion System (TadSS) were identified in all isolates. T1SS is capable of transporting unfolded substrates, including digestive enzymes (proteases and lipases), adhesins, heme-binding proteins, and proteins with repeats-in-toxins (RTX) motifs (64). T2SS, a secretion machinery prevalent in many Gram-negative bacteria, facilitates the secretion of virulence factors such as proteases, toxins, lipases, and carbohydrate-degrading enzymes, which aid the survival of both pathogenic and environmental bacteria (65). T5SS is involved in the secretion of virulence proteins critical to pathogenesis in several bacteria, including immunoglobulin A protease of *Neisseria gonorrhoeae* (*66*), IcsA protein of *Shigella flexneri* (67), and YadA of *Yersinia enterocolitica* (*68*). TadSS is an important secretion system that supports infection, cell adherence, competence, and biofilm formation in many pathogenic bacteria, such as *P*. *aeruginosa*, *Bordetella pertussis*, *Burkholderia pseudomallei*, and *Mycobacterium tuberculosis* (69). The presence of these secretion systems in all isolates suggests that they all have the potential to be pathogenic.

Most isolates have T3SS, T4SS, T6SS and flagella. T3SS is associated with the cytotoxicity of the isolates (14). AxoU is the only known T3SS substrate that is responsible for host cell cytotoxicity in *A. xylosoxidans*. All cytotoxic isolates carried both T3SS and AxoU. T4SS is primarily involved in horizontal genetic transfer (HGT) between different microorganisms, which may contribute to pathogenesis and antibiotic resistance (15). T6SS is a multiprotein complex that uses a spring-like mechanism to deliver effector proteins into target cells (70). In *A. xylosoxidans,* T6SS plays a role in targeting competing bacteria and is crucial for the internalization of the pathogen into lung epithelial cells (15). Flagella, which are evolutionarily related to T3SS, play roles in motility, surface adherence and biofilm formation (71). Interestingly, while most isolates encode flagellar genes, only 59% of them are motile, suggesting an adaptation strategy of the isolates that are isolated over a long period of time from a patient indicating chronic carriage/infection. A study on the genotypic and phenotypic evolution of the *A.xylosoxidans* isolates collected from CF patients over time showed a decrease in swimming motility, an increase in antibiotic resistance, and enhanced biofilm formation in evolved strains (72).

Phylogenetic analysis indicates that the T3SS of *A. xylosoxidans* structurally belongs to the Ysc family of T3SS systems, which includes *Bordetella* spp. (alpha-proteobacteria), *Yersinia* spp., *P. aeruginosa*, *Aeromonas* spp., *Photorhabdus luminescens*, and *Vibrio* spp. (gamma-proteobacteria) (Figure 4). In *Bordetella spp*., *Yersinia* spp., and *Photorhabdus luminescens,* the T3SS confers resistance to the innate immune response by inhibiting phagocytosis, leading to extracellular localization of the pathogens (73–75). While *P. aeruginosa* is known as an extracellular pathogen, it can replicate intracellularly where T3SS and its effectors (mainly ExoS) play critical roles in internalization, vacuole escape, evasion of inflammasome and autophagy as well as bleb formation prior to detachment (76). Similarly, T3SS is important for *Aeromonas* and *Vibrio* to maintain its intracellular niches (77, 78). In *A. xylosoxidans,* survival within macrophages for several hours is independent of T3SS; however, cytotoxicity depends on the presence of T3SS (11, 12). AxoU is the only known T3SS substrate linked to host cell lysis (14), although other T3SS effectors may be secreted during infection, as disrupting the T3SS (e.g., *sctV*) prevents cell death, whereas disruption of *axoU* alone does not (11). Interestingly, T3SS and *axoU* are expressed shortly after infection (within 30 minutes), but are subsequently turned off (11). Whether T3SS is reactivated when the bacteria escape the vacuole and lyse the host cells remain to be tested. Additional characterization of the T3SS and other virulence systems are required to identify other effectors and their role in infection and survival.

Our clinical isolates demonstrate substantial heterogeneity in their cytotoxicity (Figure 4). *A. xylosoxidans* cytotoxicity is conducted through its T3SS with the major effector known to cause cell death being AxoU, a homologue of ExoU found in *P. aeruginosa* (79). While most isolates carry T3SS and AxoU, more than 50% of isolates were non-cytotoxic, suggesting a complex regulation of T3SS expression. Multiple sequence alignment of promoter regions of T3SS genes suggests a loss of the *sctB* promoter in both GN008 and NIH-016-2. *sctB* encodes a T3SS translocon protein required for effector injection into host cells (80); therefore, its absence is expected to reduce T3SS - dependent cytotoxicity. Recent work showed that transcription of *axoU* was upregulated in GN050, but not in GN008, shortly after phagocytosis (14). Together these observations suggest that transcriptional regulation may be one mechanism controlling T3SS expression in these isolates.

The ability of *A. xylosoxidans* to adhere to host cells and undergo phagocytosis is known to be integral to its cytotoxic activity (11, 14). Multiple virulence factors have been implicated in contributing to this phenotype, including the T6SS and the adhesin ArtA (12, 15). Consistent with previous findings, our data showed that cytotoxicity across isolates required bacterial uptake (Figure 4). There was no observable relationship between overall flagellar motility and cytotoxicity but an association with biofilm production and cytotoxicity (Figure S4). In NIH-018 series, there was an increase in cytotoxicity from isolates collected after the others (Table 2). This is interesting, as in other bacteria, the bacteria tend to get less virulent when they colonized the host for a long time (81, 82). It is possible that *in vitro* cytotoxicity is not fully correlated with *in vivo* virulence, which remains to be tested.

Biofilm production is another major aspect of the bacterial lifestyle. Production of biofilm is important in promoting bacterial resistance to environmental pressures and defending against immune cells during infection (39, 83). Biofilm formation can contribute significantly to antibiotic resistance, as the biofilm matrix provides a protective environment that limits antibiotic penetration and promotes the survival of persistent cells (84). While many methods for observing biofilm formation exist, we chose to adopt static biofilm formation as it allowed for the high throughput monitoring of our available strains. Additional methods such as flow cell and micro biofilm formation are available and provide additional context for development and maintenance of biofilms, however as a diagnostic tool static biofilms are commonly used to monitor the overall production of biofilms in bacteria, and indeed in *A. xylosoxidans* (85). The NIH isolates produce low level of biofilm in general (Figure 6). We observed a slight increase in biofilm production in the intrahost isolates, NIH-010s, NIH-016s and NIH-018s although the difference was not significant. GN050 collected from ear effusion (14), is the only isolate that produced robust biofilm. It could be that because most of our isolates were from respiratory tract, they don’t form as much biofilm. Alternatively, biofilm was tightly regulated, and we didn’t assay the bacteria in the desired condition for biofilm formation in these particular isolates. Biofilm production was not correlated with motility or antibiotic resistance (Figure S4), although the strong biofilm former are moderately motile.

Flagellar motility in bacteria is commonly associated with increased disease severity and pathogenesis. Multiple related species such as *Bordetella* and *Pseudomonas* are known to have the expression of virulence systems such as the T3SS modulated by the expression of the flagella (45, 86). Additionally, bacteria with dysfunctional flagella are known to have reduced pathogenicity *in vivo* due to the inability to reach the site of infection (44). Bacteria isolated from the lungs are known to preferentially gain mutations in the flagella resulting in non-motile genotypes as compared to other sites of infection (37, 46). We identified a mutation in *flhF* which we hypothesize could result in a loss of flagellar production. We observed no correlation between overall motility and cytotoxicity (Figure S4). Interestingly, isolates that are the most cytotoxic are motile, but the most motile isolates are not cytotoxic. It could be possible that the most motile isolates can escape phagocytosis in our *in vitro* infection model. Strains that experience a loss of either flagellar motility or cytotoxicity do not experience a relative loss in the other. It is unclear what role the flagella of *A. xylosoxidans* plays during infection. In multiple bacterial species the function of the flagella is highly correlated with the activity of virulence mechanisms. The regulatory relationship between motility and virulence varies substantially between organisms. *Bordetella spp.* display an inverse relationship (86), whereas *Pseudomonas aeruginosa* displays a positive relationship between T3SS production and flagellar motility (45). *Salmonella spp.* also demonstrate a positive relationship with the SPI-1 T3SS but not the SPI-2 T3SS (87). These observations indicate a general positive correlation between flagella and virulence factor expression in extracellular pathogens. In *A. xylosoxidans,* T3SS expression seemed to increase shortly after the bacteria are phagocytosed by macrophage but dampened quickly. Whether T3SS is shut off and then gets re-activated at later stage of infection or it remains functional to some extend without making more apparatus and elevated level of substrate production requires further investigations.

In summary, this study highlights the genetic and phenotypic diversity among *A. xylosoxidans* isolates, as evidenced by the high accessory genome content and variability in secretion systems. Significant differences in antibiotic resistance and virulence mechanisms underscore the adaptability of this bacterial species. These findings emphasize the need for further mechanistic studies to elucidate the pathogenicity and antibiotic resistance mechanisms of *A. xylosoxidans*.

## 7. Methods

### Data used in this work

Raw sequence data from Bio Project ID PRJNA1148967, referred to as NIH data with 38 *Achromobacter xylosoxidans*, *A. denitrificans, A. ruhlandii, A. mucicolens and A. insolitus* genomes. 197 genomes of *Achromobacter xylosoxidans* from NCBI (https://www.ncbi.nlm.nih.gov/datasets/genome/?taxon=85698%assembly_level=1:3) as of September 2024, referred as NCBI data.

### Sources of Bacterial Strains

All NIH strains used in phenotypic studies were obtained from the NIH Clinical Center and were published in a recent work (29). GN050 and GN008 were provided by Dara Frank and have been previously published (14). PA01 Δ*wspF*Δ*pelA*Δ*pslBCD* and PA01 Δ*wspF* were previously published (88).

### Genome assembly

For all NIH isolates: Adapters were trimmed using Trimmomatic version V0.39 (89) with default parameters, except the last 10 bases were removed. FastQC (v0.11.9) (90) was used to visualize read quality, which showed an average base sequence quality greater than 28 across the reads. Genome assembly was performed using SPAdes (91) in careful mode. The quality of the assemblies was evaluated using QUAST (v5.2.0) (92), which provided metrics such as number of contigs, length, and N50.

### Genome filtering

For NIH data, contigs shorter than 500 base pairs or with coverage below 20x were removed. For NCBI data, genomes with more than 300 contigs were excluded, and contigs shorter than 500 base pairs were removed. Potential sample contamination was assessed using CheckM (v1.2.3) (93). Genomes with contamination levels exceeding 5% or completeness below 90% were excluded from subsequent analyses.

### Average Nucleotide Identity (ANI)

The complete genome sequences of *A. xylosoxidans* strains GCA_013282235.1 (GN050) and GCA_013282255.1 (GN008) were included for comparison. ANI analysis was performed using the ANIm method described previously (94) and implemented in the Python module pyANI (https://github.com/widdowquinn/pyani). ANI values were calculated for all 24 NIH genomes and the two reference strains using pyANI version 0.3.0-alpha with MUMmer-based alignments and graphical output enabled. ANI percentage identity was used to define the different clusters.

### Core genome analysis

Assembled genomes were annotated with Prokka (1.14.6) (95) with the annotation from assembly ASM1672882v1 (strain FDAARGOS_1091) used as a reference database. Core genome analysis was performed using the Panaroo pipeline version 1.5.0 (96) with a protein sequence identity threshold of 90%, gene length coverage cutoff of 75%, and a core genome sample threshold of 0.95.

### Initial phylogenetic analysis of *Achromobacter xylosoxidans*

Phylogenetic analysis of NIH and NCBI data was performed using kSNP4 (97). The optimal k-mer size was determined using Kchooser. kSNP4 was run with the selected k-mer size, and a parsimony tree was generated based on SNP loci present in at least 95% of the genomes. For NIH data, in addition to kSNP4, RaxML (98) was used to construct the phylogenetic tree. Briefly, core genes identified by Panaroo were used as input. RAxML v8.2.12 (98) was utilized to build a rooted maximum likelihood tree using the general time-reversible (GTR) substitution model with gamma correction to account for rate variation among sites. Since the phylogenetic trees generated by kSNP4 and RAxML were identical, only one tree was reported. The final tree was visualized using iTOL (99).

### Phylogenetic analysis with SNPs

To determine the phylogenetic relationships among the *A.xylosoxidans* isolates, the core genome alignment generated by Panaroo was used to identify core genome SNPs with SNP-sites (100). SNP alignment was then used to construct a maximum likelihood phylogenetic tree with IQ-TREE version 2.4 (101), employing ModelFinder Plus for model selection, ultrafast bootstrap analysis with 1000 replicates, and the approximate likelihood ratio test with 1000 replicates. The final tree was visualized and annotated using iTOL (https://itol.embl.de/).

### Population structure analysis

The genome-wide haplotype data were calculated as described previously (102). SNP calling was performed for core genes identified by the panaroo pipeline, and imputation for polymorphic sites with a missing frequency of less than <1% was conducted using BEAGLE v.3.3.2 (103). This genome-wide haplotype dataset contained 144,290 SNPs, which was used to define isolate populations and subpopulations based on haplotype profile similarity. fineSTRUCTURE analysis (104) was performed with 200,000 iterations of both the burn-in and Markov chain Monte Carlo (MCMC) method to cluster individuals based on the coancestry matrix, as previously described (105). The results were visualized as a heat map, with each cell representing the proportion of DNA a recipient receives from each donor.

### Variant calling

The variants were called with Snippy v4.4.0 (https://github.com/tseemann/snippy), with default option. NIH-016-2, collected first in the NIH-016 series, was used as a reference.

### Secretion system analysis

Secretion systems and appendages were identified using MacSyFinder 2.1.3 (106). TXSScan was run to search for all types of secretion systems and appendages, including T1SS, T2SS, T3SS, T4SS, T5SS, T6SS, Flagellum, Tad pilus, and T9SS. CONJScan was run to detect conjugative and mobilizable elements, which include T4SSs. The “ordered replicon” option was used for our complete genome sequences.

For genomes in which the secretion systems were not found by MacSyFinder, we manually looked up for key genes of the systems in the annotated genomes, as it is possible that the segmentation of the sequences that feed into MacSyFinder interferes with the gene search. These are also key genes used by MacSyFinder. A system was noted as absent when none of the genes were found.

### Phylogenetic analysis of T3SS

Amino acid sequences of SctN or its homologs were extracted from Uniprot and NCBI. Multiple sequence alignment of SctN sequences was performed using MAFFT (107) with default parameters. A rooted maximum likelihood tree was constructed using RAxML v8.2.12 (98) with the general time-reversible (GTR) substitution model and gamma correction for rate variation among sites. The phylogenetic tree was visualized using iTOL (99).

### Structural Modeling of YscN ATPase

The amino acid sequence of the T3SS ATPase from *A. xylosoxidans* was obtained from NCBI with sequence ID of WP_020925211.1. Structure of YscN was predicted using AlphaFold via the website: https://alphafoldserver.com/. The predicted structure of *Pseudomonas aeruginosa* PA14 was downloaded from https://alphafold.ebi.ac.uk/entry/A0A2U2Y0I4 with accession number of AF-A0A0H2Z9F1-F1. Structural alignment of the two proteins was performed using Pymol (The PyMOL Molecular Graphics System, Version 3.0 Schrödinger, LLC).

### Antibiotic susceptibility testing

Broth microdilution testing was carried out using two commercial Sensititre™ panels: the Gram-Negative Non-Fermenters MIC Plate and the Gram Negative MDRGN2F AST Plate (Thermo Fisher Scientific). Each isolate was adjusted to a 0.5 McFarland standard in Sensititre demineralized water and then transferred into Sensititre™ Mueller–Hinton Broth with TES. The panels were inoculated using the Sensititre™ AIM™ automated inoculation system and incubated in the Sensititre™ ARIS HiQ™ instrument according to the manufacturer’s instructions. After incubation, plates were automatically read by the system and then manually confirmed using the Sensititre™ Vizion™ Digital MIC Viewer. Quality control strains included *E. coli* ATCC 25922 and ATCC 35218, *P. aeruginosa* ATCC 27853, and *K. pneumoniae* ATCC 700603. MIC values were interpreted using CLSI breakpoints for Other Non-Enterobacterales (CLSI M100, 33rd edition, 2023).

### Biofilm

Bacterial cultures were initiated in glass culture tubes by inoculating bacteria from LB agar plates into 1 mL of LB broth at a starting OD600 of 0.1. The cultures were then incubated statically at 37°C for 3 days. On the third day, media was removed by pipetting, and biofilms were washed with water and dried overnight at 37°C. The following day, a 1% crystal violet solution was used to stain biofilms for 5 minutes, followed by additional washes with water and overnight drying at 37°C. Biofilms were solubilized with 100% DMSO, and absorbance was measured at 595 nm using a BioTek plate reader.

### Infection of *Achromobacter* in cell lines

J774a.1 cells were seeded at a density of 6×10^4^ cells in 100 μL DMEM media in a tissue culture-treated black flat-clear bottomed 96-well plate. On the day of infection, cells were washed with PBS and incubated for one hour with OPTI-MEM media containing 200 ng/mL propidium iodide (PI) and 1.45μM Hoechst 33342. *Achromobacter* inoculum was then prepared by resuspending bacteria in OPTI-MEM. Cells were infected with the bacteria at multiplicity of infection of 7 and spun down at 750 rpm for 5 minutes to synchronize infection. The 96-well plate was then inserted into a stage incubator supplied with CO_2_ and maintained at 37°C in a Nikon AXR confocal fluorescent microscope. Three images of cells were taken at arbitrary locations within each well and observed every hour for up to five hours. Cytotoxicity was measured by the ratio of PI^+^ signal to Hoechst^+^ signal (dead cells/total cells) expressed in percentage using ImageJ v 1.54g. PI⁺ and Hoechst⁺ cells were processed using background subtraction, followed by gamma adjustment and median filtering. Then, Otsu method was applied for thresholding (108), and cell counts were obtained using the Analyze Particles function.

### Adhesion and Internalization assays

J774a.1 cells were put down onto a cell culture treated 96 well plate at 6×10^4^cells/well in 100μL DMEM to grow overnight. At the day of infection, the media from the wells was replaced with media containing 1μL DMSO/mL DMEM or 2μg/mL cytochalasin D as a negative control. After 1 hour of incubation bacterial strains grown at 37^0^C then room temperature for one day each were inoculated onto the cells at an MOI of 7. After 1 hour of incubation at 37^0^C the media was removed, and the cells were washed twice with 100μL PBS. Cells which were treated with 1μL DMSO/mL DMEM had media replaced with either 1μL DMSO/mL DMEM (adherence) or with 1μL DMSO+50μg Polymyxin B/mL DMEM (internalized bacteria). Cells treated with 2μg/mL cytochalasin D had media replaced with 2μg cytochalasin D + 50μg Polymyxin B /mL (negative control). After one hour of incubation at 37^0^C the media was removed and 100μL 0.1% SDS was added and incubated at room temperature for 15 minutes. The resulting bacterial suspension was serially diluted and CFUs were measured by spot plating 5μL of dilutions onto LBA.

### Swimming motility in *Achromobacter* strains

Bacteria were inoculated and grown overnight in 1mL LB broth from single colonies on LB agar plates. Overnight growth was measured by OD600 and concentrated to an OD600 of 10 in 100μL. 2μL of the resulting suspension was injected via pipette into 0.25% agarose 2% tryptone plates. Plates were incubated at 37°C for 4 hours and imaged with a Chemidoc^tm^ Imaging System using the stain free blot procedure with 3 second exposure time. The resulting images were exported to ImageJ. Images were converted to an 8-bit format, and a brightness/contrast correction step was performed. The zone of motility was then selected using the circle tool and resulting area in Pixels was exported. Pixel count was converted to mm^2^ using the formula: mm^2^ = (Pixels)/(5000 Pixels /((3.5mm/2)^2^*3.14159)).

### Statistical analysis

All statistical analyses were performed in GraphPad Prism version 10. Data were assessed for normality using the Shapiro-Wilk test. For comparisons among multiple groups, one-way ANOVA with Dunnett’s post hoc test was used for normally distributed data, while the nonparametric Kruskal-Wallis test followed by Dunn’s multiple comparisons test was used for nonparametric data. Data are presented as mean ± SD unless otherwise stated.

## 8. Author statements

### 8.1 Author contributions

Pooja Acharya (Formal Analysis, Data Curation, Methodology, Writing), Cameron Lloyd (Formal Analysis, Methodology, Writing), Ngoc Thien Lam (Formal Analysis, Writing), Jessica Kumke (Formal Analysis), Sreejana Ray (Resources), Zilia Yanira Muñoz Ramirez (Formal Analysis), Sanchita Das (conceptualization, resources) and Hanh Ngoc Lam (Conceptualization, Data Curation, Formal Analysis, Funding Acquisition, Investigation, Methodology, Supervision, Writing)

## 8.4 Acknowledgements

We thank Dara Frank (medical College of Wisconsin) for sharing the GN050 and GN008 strains.

## References

1. Veschetti L, Boaretti M, Saitta GM, Passarelli Mantovani R, Lleo MM, Sandri A, et al. Achromobacter spp. prevalence and adaptation in cystic fibrosis lung infection. Microbiol Res. 2022;263:127140.

2. Jeukens J, Freschi L, Vincent AT, Emond-Rheault JG, Kukavica-Ibrulj I, Charette SJ, et al. A Pan-Genomic Approach to Understand the Basis of Host Adaptation in Achromobacter. Genome Biol Evol. 2017;9(4):1030–46.

3. Yabuuchi E, Oyama A. Achromobacter xylosoxidans n. sp. from human ear discharge. Jpn J Microbiol. 1971;15(5):477–81.

4. Swenson CE, Sadikot RT. Achromobacter respiratory infections. Ann Am Thorac Soc. 2015;12:252–8.

5. Jakobsen TH, Hansen MA, Jensen PØ, Hansen L, Riber L, Cockburn A, et al. Complete Genome Sequence of the Cystic Fibrosis Pathogen Achromobacter xylosoxidans NH44784-1996 Complies with Important Pathogenic Phenotypes. PLoS ONE. 2013;8(7):e68484.

6. Esposito S, Pisi G, Fainardi V, Principi N. What is the role of Achromobacter species in patients with cystic fibrosis? Front Biosci (Landmark Ed). 2021;26(12):1613–20.

7. Tetart M, Wallet F, Kyheng M, Leroy S, Perez T, Le Rouzic O, et al. Impact of Achromobacter xylosoxidans isolation on the respiratory function of adult patients with cystic fibrosis. ERJ Open Res. 2019;5(4).

8. Somayaji R, Stanojevic S, Tullis DE, Stephenson AL, Ratjen F, Waters V. Clinical Outcomes Associated with Achromobacter Species Infection in Patients with Cystic Fibrosis. Ann Am Thorac Soc. 2017;14(9):1412–8.

9. Nielsen SM, Penstoft LN, Nørskov-Lauritsen N. Motility, Biofilm Formation and Antimicrobial Efflux of Sessile and Planktonic Cells of Achromobacter xylosoxidans. Pathogens. 2019;8(1):14.

10. Wills BM, Garai P, Riegert MO, Sanchez FT, Pickrum AM, Frank DW, et al. Identification of Virulence Factors Involved in a Murine Model of Severe Achromobacter xylosoxidans Infection. Infect Immun. 2023;91(7):e0003723.

11. Turton K, Parks HJ, Zarodkiewicz P, Hamad MA, Dwane R, Parau G, et al. The Achromobacter type 3 secretion system drives pyroptosis and immunopathology via independent activation of NLRC4 and NLRP3 inflammasomes. Cell Rep. 2023;42(8):113012.

12. Pickrum AM, Riegert MO, Wells C, Brockman K, Frank DW. The In Vitro Replication Cycle of Achromobacter xylosoxidans and Identification of Virulence Genes Associated with Cytotoxicity in Macrophages. Microbiol Spectr. 2022;10(4):e0208322.

13. Li X, Hu Y, Gong J, Zhang L, Wang G. Comparative genome characterization of Achromobacter members reveals potential genetic determinants facilitating the adaptation to a pathogenic lifestyle. Appl Microbiol Biotechnol. 2013;97(14):6413–25.

14. Pickrum AM, DeLeon O, Dirck A, Tessmer MH, Riegert MO, Biller JA, et al. Achromobacter xylosoxidans Cellular Pathology Is Correlated with Activation of a Type III Secretion System. Infect Immun. 2020;88(7).

15. Le Goff M, Vastel M, Lebrun R, Mansuelle P, Diarra A, Grandjean T, et al. Characterization of the Achromobacter xylosoxidans Type VI Secretion System and Its Implication in Cystic Fibrosis. Front Cell Infect Microbiol. 2022;12:859181.

16. Troisfontaines P, Cornelis GR. Type III secretion: more systems than you think. Physiology (Bethesda). 2005;20:326–39.

17. Weyrich LS, Rolin OY, Muse SJ, Park J, Spidale N, Kennett MJ, et al. A Type VI secretion system encoding locus is required for Bordetella bronchiseptica immunomodulation and persistence in vivo. PLoS One. 2012;7(10):e45892.

18. Nolan LM, Allsopp LP. Antimicrobial Weapons of Pseudomonas aeruginosa. Adv Exp Med Biol. 2022;1386:223–56.

19. Li X, Hu Y, Gong J, Zhang L, Wang G. Comparative genome characterization of Achromobacter members reveals potential genetic determinants facilitating the adaptation to a pathogenic lifestyle. Applied Microbiology and Biotechnology. 2013;97(14):6413–25.

20. Aggarwal S, Garcia-Telles N, Aggarwal G, Lavie C, Lippi G, Henry BM. Clinical features, laboratory characteristics, and outcomes of patients hospitalized with coronavirus disease 2019 (COVID-19): Early report from the United States. Diagnosis (Berl). 2020;7(2):91–6.

21. Lorenz A, Preusse M, Bruchmann S, Pawar V, Grahl N, Pils MC, et al. Importance of flagella in acute and chronic Pseudomonas aeruginosa infections. Environ Microbiol. 2019;21(3):883–97.

22. Isler B, Kidd TJ, Stewart AG, Harris P, Paterson DL. Achromobacter Infections and Treatment Options. Antimicrob Agents Chemother. 2020;64(11).

23. Swenson CE, Sadikot RT. Achromobacter respiratory infections. Ann Am Thorac Soc. 2015;12(2):252–8.

24. Hu Y, Zhu Y, Ma Y, Liu F, Lu N, Yang X, et al. Genomic insights into intrinsic and acquired drug resistance mechanisms in Achromobacter xylosoxidans. Antimicrob Agents Chemother. 2015;59(2):1152–61.

25. Khaledi A, Weimann A, Schniederjans M, Asgari E, Kuo TH, Oliver A, et al. Predicting antimicrobial resistance in Pseudomonas aeruginosa with machine learning-enabled molecular diagnostics. EMBO Mol Med. 2020;12(3):e10264.

26. Hyun JC, Kavvas ES, Monk JM, Palsson BO. Machine learning with random subspace ensembles identifies antimicrobial resistance determinants from pan-genomes of three pathogens. PLoS Comput Biol. 2020;16(3):e1007608.

27. Kavvas ES, Catoiu E, Mih N, Yurkovich JT, Seif Y, Dillon N, et al. Machine learning and structural analysis of Mycobacterium tuberculosis pan-genome identifies genetic signatures of antibiotic resistance. Nat Commun. 2018;9(1):4306.

28. Nguyen M, Long SW, McDermott PF, Olsen RJ, Olson R, Stevens RL, et al. Using Machine Learning To Predict Antimicrobial MICs and Associated Genomic Features for Nontyphoidal Salmonella. J Clin Microbiol. 2019;57(2).

29. Ray S, Flemming LK, Scudder CJ, Ly MA, Porterfield HS, Smith RD, et al. Comparative phenotypic and genotypic antimicrobial susceptibility surveillance in Achromobacter spp. through whole genome sequencing. Microbiol Spectr. 2025;13(4):e0252724.

30. Meier-Kolthoff JP, Carbasse JS, Peinado-Olarte RL, Goker M. TYGS and LPSN: a database tandem for fast and reliable genome-based classification and nomenclature of prokaryotes. Nucleic Acids Res. 2022;50(D1):D801–D7.

31. Kalantar KL, Carvalho T, de Bourcy CFA, Dimitrov B, Dingle G, Egger R, et al. IDseq-An open source cloud-based pipeline and analysis service for metagenomic pathogen detection and monitoring. Gigascience. 2020;9(10).

32. Bador J, Neuwirth C, Grangier N, Muniz M, Germe L, Bonnet J, et al. Role of AxyZ Transcriptional Regulator in Overproduction of AxyXY-OprZ Multidrug Efflux System in Achromobacter Species Mutants Selected by Tobramycin. Antimicrob Agents Chemother. 2017;61(8).

33. Rees VE, Deveson Lucas DS, Lopez-Causape C, Huang Y, Kotsimbos T, Bulitta JB, et al. Characterization of Hypermutator Pseudomonas aeruginosa Isolates from Patients with Cystic Fibrosis in Australia. Antimicrob Agents Chemother. 2019;63(4).

34. Ragheb MN, Thomason MK, Hsu C, Nugent P, Gage J, Samadpour AN, et al. Inhibiting the Evolution of Antibiotic Resistance. Mol Cell. 2019;73(1):157–65 e5.

35. Davidson AL, Chen J. ATP-binding cassette transporters in bacteria. Annu Rev Biochem. 2004;73:241–68.

36. Chen L, Zou Y, She P, Wu Y. Composition, function, and regulation of T6SS in Pseudomonas aeruginosa. Microbiol Res. 2015;172:19–25.

37. Holban AM, Gregoire CM, Gestal MC. Conquering the host: Bordetella spp. and Pseudomonas aeruginosa molecular regulators in lung infection. Front Microbiol. 2022;13:983149.

38. Warrier A, Satyamoorthy K, Murali TS. Quorum-sensing regulation of virulence factors in bacterial biofilm. Future Microbiol. 2021;16:1003–21.

39. Yin R, Cheng J, Wang J, Li P, Lin J. Treatment of Pseudomonas aeruginosa infectious biofilms: Challenges and strategies. Front Microbiol. 2022;13:955286.

40. Nielsen SM, Penstoft LN, Norskov-Lauritsen N. Motility, Biofilm Formation and Antimicrobial Efflux of Sessile and Planktonic Cells of Achromobacter xylosoxidans. Pathogens. 2019;8(1).

41. Passos da Silva D, Matwichuk ML, Townsend DO, Reichhardt C, Lamba D, Wozniak DJ, et al. The Pseudomonas aeruginosa lectin LecB binds to the exopolysaccharide Psl and stabilizes the biofilm matrix. Nat Commun. 2019;10(1):2183.

42. Starkey M, Hickman JH, Ma L, Zhang N, De Long S, Hinz A, et al. Pseudomonas aeruginosa rugose small-colony variants have adaptations that likely promote persistence in the cystic fibrosis lung. J Bacteriol. 2009;191(11):3492–503.

43. Kazmierczak BI, Schniederberend M, Jain R. Cross-regulation of Pseudomonas motility systems: the intimate relationship between flagella, pili and virulence. Curr Opin Microbiol. 2015;28:78–82.

44. Feldman M, Bryan R, Rajan S, Scheffler L, Brunnert S, Tang H, et al. Role of flagella in pathogenesis of Pseudomonas aeruginosa pulmonary infection. Infect Immun. 1998;66(1):43–51.

45. Soscia C, Hachani A, Bernadac A, Filloux A, Bleves S. Cross talk between type III secretion and flagellar assembly systems in Pseudomonas aeruginosa. J Bacteriol. 2007;189(8):3124–32.

46. Planet PJ. Adaptation and Evolution of Pathogens in the Cystic Fibrosis Lung. J Pediatric Infect Dis Soc. 2022;11(Supplement_2):S23–S31.

47. Schniederberend M, Abdurachim K, Murray TS, Kazmierczak BI. The GTPase activity of FlhF is dispensable for flagellar localization, but not motility, in Pseudomonas aeruginosa. J Bacteriol. 2013;195(5):1051–60.

48. Lloyd CJ, Klose KE. The Vibrio Polar Flagellum: Structure and Regulation. Adv Exp Med Biol. 2023;1404:77–97.

49. Blum M, Andreeva A, Florentino LC, Chuguransky SR, Grego T, Hobbs E, et al. InterPro: the protein sequence classification resource in 2025. Nucleic Acids Res. 2025;53(D1):D444–D56.

50. Bange G, Petzold G, Wild K, Parlitz RO, Sinning I. The crystal structure of the third signal-recognition particle GTPase FlhF reveals a homodimer with bound GTP. Proc Natl Acad Sci U S A. 2007;104(34):13621–5.

51. Zhang K, He J, Cantalano C, Guo Y, Liu J, Li C. FlhF regulates the number and configuration of periplasmic flagella in Borrelia burgdorferi. Mol Microbiol. 2020;113(6):1122–39.

52. Balaban M, Joslin SN, Hendrixson DR. FlhF and its GTPase activity are required for distinct processes in flagellar gene regulation and biosynthesis in Campylobacter jejuni. J Bacteriol. 2009;191(21):6602–11.

53. Jumper J, Evans R, Pritzel A, Green T, Figurnov M, Ronneberger O, et al. Highly accurate protein structure prediction with AlphaFold. Nature. 2021;596(7873):583–9.

54. Papalia M, Figueroa-Espinosa R, Steffanowski C, Barberis C, Almuzara M, Barrios R, et al. Expansion and improvement of MALDI-TOF MS databases for accurate identification of Achromobacter species. J Microbiol Methods. 2020;172:105889.

55. Mosquera-Rendon J, Rada-Bravo AM, Cardenas-Brito S, Corredor M, Restrepo-Pineda E, Benitez-Paez A. Pangenome-wide and molecular evolution analyses of the Pseudomonas aeruginosa species. Bmc Genomics. 2016;17:45.

56. Abram KZ, Jun SR, Udaondo Z. Pseudomonas aeruginosa Pangenome: Core and Accessory Genes of a Highly Resourceful Opportunistic Pathogen. Adv Exp Med Biol. 2022;1386:3–28.

57. Thorell K, Muñoz-Ramírez ZY, Wang D, Sandoval-Motta S, Boscolo Agostini R, Ghirotto S, et al. The Helicobacter pylori Genome Project: insights into H. pylori population structure from analysis of a worldwide collection of complete genomes. Nature Communications. 2023;14(1).

58. Salama NR, Shepherd B, Falkow S. Global transposon mutagenesis and essential gene analysis of Helicobacter pylori. J Bacteriol. 2004;186(23):7926–35.

59. Bador J, Amoureux L, Duez J-M, Drabowicz A, Siebor E, Llanes C, et al. First Description of an RND-Type Multidrug Efflux Pump in Achromobacter xylosoxidans, AxyABM. Antimicrobial Agents and Chemotherapy. 2011;55(10):4912–4.

60. Bador J, Amoureux L, Blanc E, Neuwirth C. Innate Aminoglycoside Resistance of Achromobacter xylosoxidans Is Due to AxyXY-OprZ, an RND-Type Multidrug Efflux Pump. Antimicrobial Agents and Chemotherapy. 2013;57(1):603–5.

61. Magallon A, Bador J, Garrigos T, Demeule C, Chapelle A, Varin V, et al. Description of Two Resistance-Nodulation-Cell Division Efflux Systems Involved in Acquired Antibiotic Resistance: AxySUV in Achromobacter xylosoxidans and AinCDJ in Achromobacter insuavis. Antibiotics (Basel). 2025;14(6).

62. Ventero MP, Haro-Moreno JM, Molina-Pardines C, Sanchez-Bautista A, Garcia-Rivera C, Boix V, et al. Role of Relebactam in the Antibiotic Resistance Acquisition in Pseudomonas aeruginosa: In Vitro Study. Antibiotics (Basel). 2023;12(11).

63. Palomba E, Comelli A, Saluzzo F, Di Marco F, Matarazzo E, Re NL, et al. Activity of imipenem/relebactam against KPC-producing Klebsiella pneumoniae and the possible role of Ompk36 mutation in determining resistance: an Italian retrospective analysis. Ann Clin Microbiol Antimicrob. 2025;24(1):23.

64. Holland IB, Schmitt L, Young J. Type 1 protein secretion in bacteria, the ABC-transporter dependent pathway (review). Mol Membr Biol. 2005;22(1-2):29–39.

65. Korotkov KV, Sandkvist M. Architecture, Function, and Substrates of the Type II Secretion System. EcoSal Plus. 2019;8(2):227–44.

66. Pohlner J, Halter R, Meyer TF. Neisseria gonorrhoeae IgA protease. Secretion and implications for pathogenesis. Antonie Van Leeuwenhoek. 1987;53(6):479–84.

67. Brandon LD, Goehring N, Janakiraman A, Yan AW, Wu T, Beckwith J, et al. IcsA, a polarly localized autotransporter with an atypical signal peptide, uses the Sec apparatus for secretion, although the Sec apparatus is circumferentially distributed. Mol Microbiol. 2003;50(1):45–60.

68. Roggenkamp A, Ackermann N, Jacobi CA, Truelzsch K, Hoffmann H, Heesemann J. Molecular analysis of transport and oligomerization of the Yersinia enterocolitica adhesin YadA. J Bacteriol. 2003;185(13):3735–44.

69. Tassinari M, Rudzite M, Filloux A, Low HH. Assembly mechanism of a Tad secretion system secretin-pilotin complex. Nature Communications. 2023;14(1).

70. Cherrak Y, Flaugnatti N, Durand E, Journet L, Cascales E. Structure and Activity of the Type VI Secretion System. Microbiology Spectrum. 2019;7(4):329–42.

71. Zhang X, Zeng W, Kong J, Huang Z, Shu H, Tang M, et al. The prevalence and mechanisms of heteroresistance to ceftazidime/avibactam in KPC-producing Klebsiella pneumoniae. J Antimicrob Chemother. 2024;79(8):1865–76.

72. Khademi SMH, Gabrielaite M, Paulsson M, Knulst M, Touriki E, Marvig RL, et al. Genomic and Phenotypic Evolution of Achromobacter xylosoxidans during Chronic Airway Infections of Patients with Cystic Fibrosis. mSystems. 2021;6(3).

73. Seabaugh JA, Anderson DM. Pathogenicity and virulence of Yersinia. Virulence. 2024;15(1):2316439.

74. Yuk MH, Harvill ET, Cotter PA, Miller JF. Modulation of host immune responses, induction of apoptosis and inhibition of NF-kappaB activation by the Bordetella type III secretion system. Mol Microbiol. 2000;35(5):991–1004.

75. Brugirard-Ricaud K, Duchaud E, Givaudan A, Girard PA, Kunst F, Boemare N, et al. Site-specific antiphagocytic function of the Photorhabdus luminescens type III secretion system during insect colonization. Cell Microbiol. 2005;7(3):363–71.

76. Resko ZJ, Suhi RF, Thota AV, Kroken AR. Evidence for intracellular Pseudomonas aeruginosa. J Bacteriol. 2024;206(5):e0010924.

77. de Souza Santos M, Orth K. Intracellular Vibrio parahaemolyticus escapes the vacuole and establishes a replicative niche in the cytosol of epithelial cells. mBio. 2014;5(5):e01506–14.

78. Guerra RM, Maleno FD, Figueras MJ, Pujol-Bajador I, Fernandez-Bravo A. Potential Pathogenicity of Aeromonas spp. Recovered in River Water, Soil, and Vegetation from a Natural Recreational Area. Pathogens. 2022;11(11).

79. Tessmer MH, Anderson DM, Pickrum AM, Riegert MO, Frank DW. Identification and Verification of Ubiquitin-Activated Bacterial Phospholipases. J Bacteriol. 2019;201(4).

80. Dey S, Chakravarty A, Guha Biswas P, De Guzman RN. The type III secretion system needle, tip, and translocon. Protein Sci. 2019;28(9):1582–93.

81. Vanderwoude J, Fleming D, Azimi S, Trivedi U, Rumbaugh KP, Diggle SP. The evolution of virulence in Pseudomonas aeruginosa during chronic wound infection. Proc Biol Sci. 2020;287(1937):20202272.

82. Didelot X, Walker AS, Peto TE, Crook DW, Wilson DJ. Within-host evolution of bacterial pathogens. Nat Rev Microbiol. 2016;14(3):150–62.

83. Uruen C, Chopo-Escuin G, Tommassen J, Mainar-Jaime RC, Arenas J. Biofilms as Promoters of Bacterial Antibiotic Resistance and Tolerance. Antibiotics (Basel). 2020;10(1).

84. Yan J, Bassler BL. Surviving as a Community: Antibiotic Tolerance and Persistence in Bacterial Biofilms. Cell Host Microbe. 2019;26(1):15–21.

85. Sandri A, Veschetti L, Saitta GM, Passarelli Mantovani R, Carelli M, Burlacchini G, et al. Achromobacter spp. Adaptation in Cystic Fibrosis Infection and Candidate Biomarkers of Antimicrobial Resistance. Int J Mol Sci. 2022;23(16).

86. Kurushima J, Kuwae A, Abe A. The type III secreted protein BspR regulates the virulence genes in Bordetella bronchiseptica. PLoS One. 2012;7(6):e38925.

87. Singer HM, Kuhne C, Deditius JA, Hughes KT, Erhardt M. The Salmonella Spi1 virulence regulatory protein HilD directly activates transcription of the flagellar master operon flhDC. J Bacteriol. 2014;196(7):1448–57.

88. Colvin KM, Irie Y, Tart CS, Urbano R, Whitney JC, Ryder C, et al. The Pel and Psl polysaccharides provide Pseudomonas aeruginosa structural redundancy within the biofilm matrix. Environ Microbiol. 2012;14(8):1913–28.

89. Bolger AM, Lohse M, Usadel B. Trimmomatic: a flexible trimmer for Illumina sequence data. Bioinformatics. 2014;30(15):2114–20.

90. Andrews S. FastQC: a quality control tool for high throughput sequence data. 2010.

91. Bankevich A, Nurk S, Antipov D, Gurevich AA, Dvorkin M, Kulikov AS, et al. SPAdes: a new genome assembly algorithm and its applications to single-cell sequencing. J Comput Biol. 2012;19(5):455–77.

92. Gurevich A, Saveliev V, Vyahhi N, Tesler G. QUAST: quality assessment tool for genome assemblies. Bioinformatics. 2013;29(8):1072–5.

93. Parks DH, Imelfort M, Skennerton CT, Hugenholtz P, Tyson GW. CheckM: assessing the quality of microbial genomes recovered from isolates, single cells, and metagenomes. Genome Res. 2015;25(7):1043–55.

94. Richter M, Rossello-Mora R. Shifting the genomic gold standard for the prokaryotic species definition. Proc Natl Acad Sci U S A. 2009;106(45):19126–31.

95. Seemann T. Prokka: rapid prokaryotic genome annotation. Bioinformatics. 2014;30(14):2068–9.

96. Tonkin-Hill G, MacAlasdair N, Ruis C, Weimann A, Horesh G, Lees JA, et al. Producing polished prokaryotic pangenomes with the Panaroo pipeline. Genome Biol. 2020;21(1):180.

97. Hall BG, Nisbet J. Building Phylogenetic Trees From Genome Sequences With kSNP4. Mol Biol Evol. 2023;40(11).

98. Stamatakis A. RAxML version 8: a tool for phylogenetic analysis and post-analysis of large phylogenies. Bioinformatics. 2014;30(9):1312–3.

99. Letunic I, Bork P. Interactive Tree of Life (iTOL) v6: recent updates to the phylogenetic tree display and annotation tool. Nucleic Acids Res. 2024;52(W1):W78–W82.

100. Page AJ, Taylor B, Delaney AJ, Soares J, Seemann T, Keane JA, et al. SNP-sites: rapid efficient extraction of SNPs from multi-FASTA alignments. Microb Genom. 2016;2(4):e000056.

101. Minh BQ, Schmidt HA, Chernomor O, Schrempf D, Woodhams MD, von Haeseler A, et al. IQ-TREE 2: New Models and Efficient Methods for Phylogenetic Inference in the Genomic Era. Mol Biol Evol. 2020;37(5):1530–4.

102. Yahara K, Didelot X, Ansari MA, Sheppard SK, Falush D. Efficient inference of recombination hot regions in bacterial genomes. Mol Biol Evol. 2014;31(6):1593–605.

103. Browning BL, Browning SR. A unified approach to genotype imputation and haplotype-phase inference for large data sets of trios and unrelated individuals. Am J Hum Genet. 2009;84(2):210–23.

104. Lawson DJ, Hellenthal G, Myers S, Falush D. Inference of population structure using dense haplotype data. PLoS Genet. 2012;8(1):e1002453.

105. Yahara K, Furuta Y, Oshima K, Yoshida M, Azuma T, Hattori M, et al. Chromosome painting in silico in a bacterial species reveals fine population structure. Mol Biol Evol. 2013;30(6):1454–64.

106. Abby SS, Cury J, Guglielmini J, Neron B, Touchon M, Rocha EP. Identification of protein secretion systems in bacterial genomes. Sci Rep. 2016;6:23080.

107. Katoh K, Standley DM. MAFFT multiple sequence alignment software version 7: improvements in performance and usability. Mol Biol Evol. 2013;30(4):772–80.

108. Otsu N. A Threshold Selection Method from Gray-Level Histograms. IEEE Transactions on Systems, Man, and Cybernetics. 1979;9(1):62–6.

109. Jones P, Binns D, Chang HY, Fraser M, Li W, McAnulla C, et al. InterProScan 5: genome-scale protein function classification. Bioinformatics. 2014;30(9):1236–40.

